# Selection on plastic adherence leads to hyper-multicellular strains and incidental virulence in the budding yeast

**DOI:** 10.1101/2022.06.03.494655

**Authors:** Luke I. Ekdahl, Juliana A. Salcedo, Matthew M. Dungan, Despina V. Mason, Dulguun Myagmarsuren, Helen A. Murphy

## Abstract

Many disease-causing microbes are not obligate pathogens; rather, they are environmental microbes taking advantage of an ecological opportunity. The existence of microbes that are not normally pathogenic, yet are well-suited to host exploitation, is an evolutionary paradox. One hypothesis posits that selection in the environment may favor traits that incidentally lead to pathogenicity and virulence, or serve as pre-adaptations for survival in a host. An example of such a trait is surface adherence. To experimentally test the idea of “accidental virulence”, replicate populations of the yeast, *Saccharomyces cerevisiae*, which can be an opportunistic pathogen, were evolved to attach to a plastic bead for hundreds of generations. Along with plastic adherence, two multicellular phenotypes— biofilm formation and flor formation— increased; another phenotype, pseudohyphal growth, responded to the nutrient limitation. Thus, experimental selection led to the evolution of highly-adherent, hyper-multicellular strains. Wax moth larvae injected with evolved hyper-multicellular strains were significantly more likely to die than those injected with evolved non-multicellular strains. Hence, selection on plastic adherence incidentally led to the evolution of enhanced multicellularity and increased virulence. Our results support the idea that selection in the environment for a trait unrelated to virulence can inadvertently generate opportunistic, “accidental” pathogens.

## Introduction

The study of infectious disease often focuses on pathogenic microbes that either specialize on exploiting animal hosts or on commensals that switch to pathogenesis when the delicate balance between host and microbe is perturbed. These microbes are presumed to have co-evolved complex adaptations that allow survival and reproduction in and on hosts. However, there exists a broad range of microbial organisms that live in the open environment (i.e., soil, vegetation, aquatic habitats) that are capable of causing disease when the opportunity presents itself [1]. Such microbes also have adaptations that allow host exploitation, but the origin of these adaptations is unclear, as growth and survival in a host is not a required part of the lifecycle [2]. For example, the soil-associated bacteria *Pseudomonas aeruginosa* [3] and *Burkholderia cepacia* [4] can infect the lungs of cystic fibrosis patients.

The ‘accidental virulence’ hypothesis [2] proposes that adapting to harsh environmental conditions can favor traits that allow certain microbes to be well-suited to exploiting hosts, and challenges the idea that intricate co-evolution is a requirement of microbial pathogenesis and virulence. For example, the fungus *Cryptococcus neoformans* may have ‘dual use’ virulence factors, such as capsule formation, that are favored in the environment, but that also provide advantages in animal hosts [5]. Other examples include thermotolerance and halotolerance, which may make colonization in and on humans more likely [6,7]. The yeast *Candida auris*, which is found in coastal wetlands, can cause severe systemic infection [8]. One last example is adherence, which is important for many microbial behaviors required for survival (e.g., biofilm formation) [9], but that can also play a role in pathogenicity and virulence [10,11]. In the soil-associated yeast, *Blastomyces dermatitidis* [12], which can cause lung infections, the deletion of a single adhesin gene abolishes pathogenesis [13]. While the function of microbial traits in both the open environment and the host are crucial evidence for the accidental virulence hypothesis, the hypothesis has not been directly tested experimentally.

To test accidental virulence, we evolved populations of the biomedical model yeast, *Saccharomyces cerevisiae*. Aside from being found in a myriad of ecological niches around the globe [14], *S. cerevisiae* is an opportunistic pathogen capable of infecting immunocompromised individuals, with reports of infections increasing [15–20]. As such, it has been a model for fungal pathogenesis-related traits [21–24]. Some environmentally-derived strains are capable of adhering to surfaces and expressing associated aggregative phenotypes [25]. These range from biofilms on solid and semi-solid agar, to invasive and pseudohyphal growth, to floating mats on liquid surfaces (flors). Of these multicellular phenotypes, only invasive and pseudohyphal growth have been linked to pathogenicity in *S. cerevisiae* [23,24,26], although biofilm formation has been linked to pathogenicity in other fungi [27]. Not all strains are capable of expressing these phenotypes; and while some strains express multiple multicellular traits [28,29], there does not appear to be a correlation among the numerous phenotypes [30]. The ability of a strain to express multicellularity in one form does not necessarily suggest the ability to express it in another, despite overlap in conserved signaling and regulatory networks governing the traits [31]. This is not entirely surprising, since different environmental conditions likely favor different multicellular phenotypes.

Here, yeast populations were artificially selected for adherence ability in one context, in order to determine whether it led to an increase in virulence in another. Specifically, yeast were evolved to adhere to a plastic bead [32], then tested against wax moth larvae to estimate virulence. In replicate populations of two genetic backgrounds, the yeast increased in their ability to express multicellularity in numerous forms. Not all multicellular phenotypes responded in the same way, with pseudohyphal growth appearing to evolve independently from plastic adherence, biofilm formation and flor formation. This phenotypic evolution demonstrates the complexity of the interacting genetic networks underlying yeast multicellularity. Along with these correlated effects of selection, the yeast also became more virulent. Our results experimentally demonstrate that selection outside of a host environment can inadvertently favor traits that serve as preadaptations for virulence.

## Results

The evolution experiment was conducted in two genetic backgrounds. While most *S. cerevisiae* strains can be pathogenic against wax moth larvae if administered with a high enough inoculum [24], we chose two strains isolated from clinical settings, and therefore, had a known tendency toward human pathogenicity as well. These strains were highly heterozygous, with tens of thousands of SNPs in each genome [33]; thus, the strains contained standing genetic variation on which selection could act. The first strain, YJM311, was isolated from the bile tube of a patient in San Francisco in 1981 [34]; its recombinant offspring vary in at least one form of filamentous growth [35]. The second strain, YJM128, was isolated from the lung of a patient in Missouri in the 1980’s [36]. Both strains were engineered to constitutively express mCherry, then sporulated, digested, and germinated. Each pool of recombinant offspring was used to inoculate replicate populations.

From each ancestral strain, ten replicate populations were evolved via serial transfer for 350-400 mitotic generations, half punctuated with sexual cycles every 40 generations, and two without beads as controls. YJM311 was evolved for 8 sexual cycles, while YJM128 was evolved for 9. Populations were grown in limiting medium in glass tubes in the presence of a plastic bead (Figure 1A-B). After growth, beads were washed, suspended in water, and sonicated to detach cells. The cell suspension was transferred to the next tube for growth (Figure S1). In YJM128, the sexual control failed to propagate after the first cycle. Therefore, a full complement of controls (3 asexual and 3 sexual) were subsequently initiated and evolved in the same manner.

**Figure 1:**
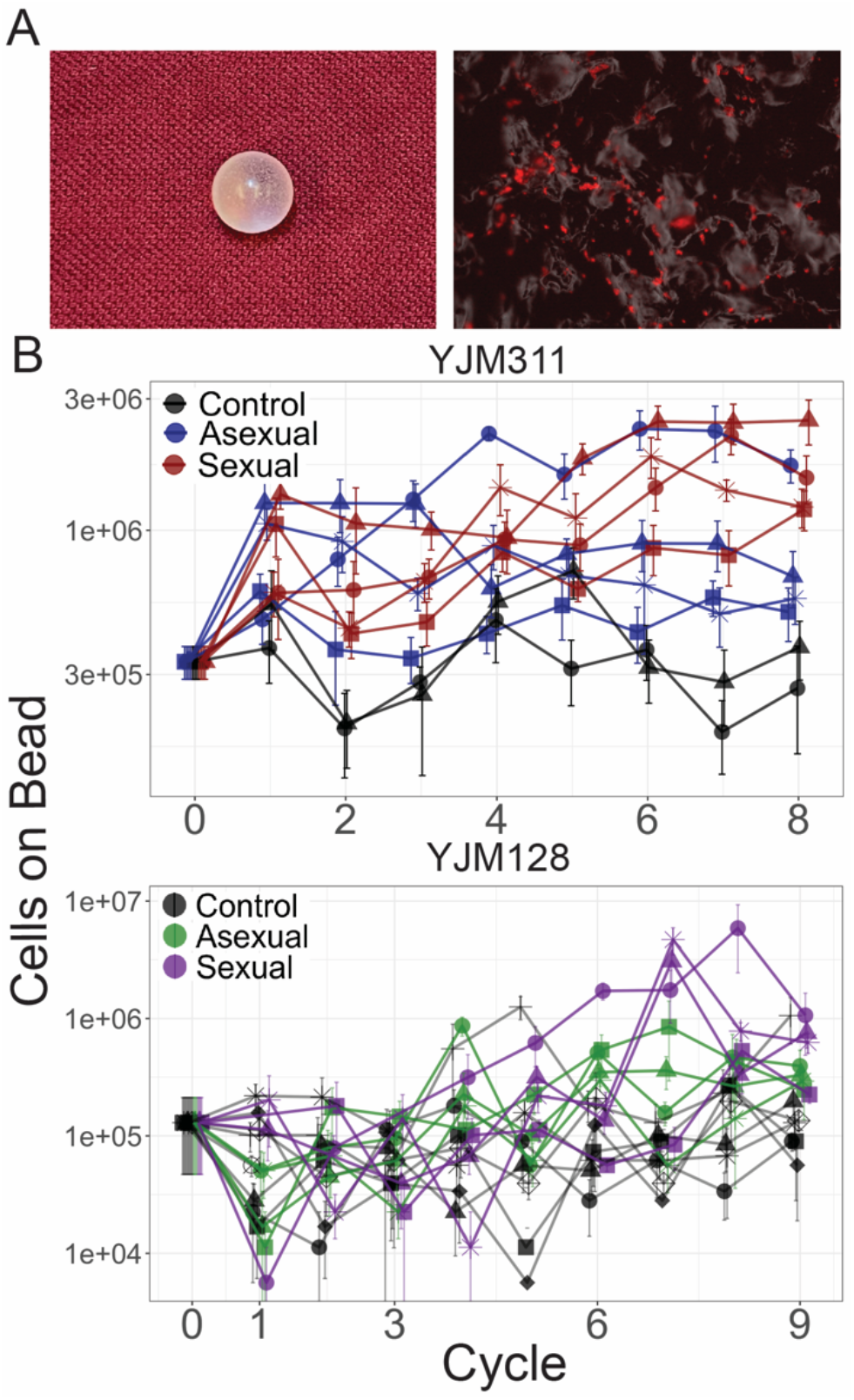
Evolution of bead adherence. **(A)** Image of a 7mm experimental bead; close-up image with attached cells expressing mCherry. **(B)** Whole population-bead adherence of replicate populations over the experimental cycles. Y-axis plots the number of cells adhering to a plastic bead, as estimated by hemocytometer counts (± S.E.M.) on a log scale. Along with the ancestral timepoint, for each population at each timepoint, cells from 8 replicate beads were counted in YJM311 (670 beads in total); for YJM128, 4 replicate beads were counted (542 beads in total).

The number of cells attaching to the bead increased over time in the experimental populations (Figure 1; Table S3); thus, the experimental selection protocol had the desired effect. As has been demonstrated in other evolution experiments in microorganisms [37–42], sexual populations showed increased adaptation in comparison to asexual populations (linear mixed-effect model (LMM) with log-transformed bead cell counts: YJM311, asexual*cycle coefficient = 0.054 (confidence interval ±0.048), sexual*cycle = 0.184 (±0.048); YJM128, asexual*cycle = 0.47 (±0.242), sexual*cycle = 0.598 (±0.254)).

At the end of the experiment, ten individual clones were isolated from each population from four timepoints (for YJM311, cycles 2, 4, 6, 8, and for YJM128, cycles 1, 3, 6, 9). For each genetic background, over 400 clones, along with 20 ancestral recombinant offspring, were arrayed in a 96-well plate format for analysis of multicellular phenotypes.

### Plastic Adherence Ability

The panel of clones was first assayed for plastic adherence ability (Figure 2A). Plastic adherence was measured with a microplate reader that detected the fluorescent signal of cells remaining attached to a well in which culture was grown to saturation and gently rinsed. As expected, plastic adherence increased over time in the clones from experimental populations (LMM: YJM311, control*cycle coefficient = 0.008 (confidence interval ±0.056), asexual*cycle = 0.040 (±0.040), sexual*cycle = 0.100 (±0.040); YJM128, control*cycle= 0.016 (±0.076), asexual*cycle = 0.012 (±0.102); sexual*cycle = 0.187 (±0.114); Table S4). Fluorescent signal could have evolved over the course of the experiment; indeed, from the beginning, YJM128 produced a brighter fluorescent signal than YJM311, suggesting the existence of genetic variants that could influence fluorescence expression. Despite the potential for noise in the measurement, the signal of increased adherence throughout the experiment was apparent in both genetic backgrounds. These clonal data support the results of the whole-population adherence measurement (Figure 1), in which cells attaching to a plastic bead were counted manually with a hemocytometer.

**Figure 2:**
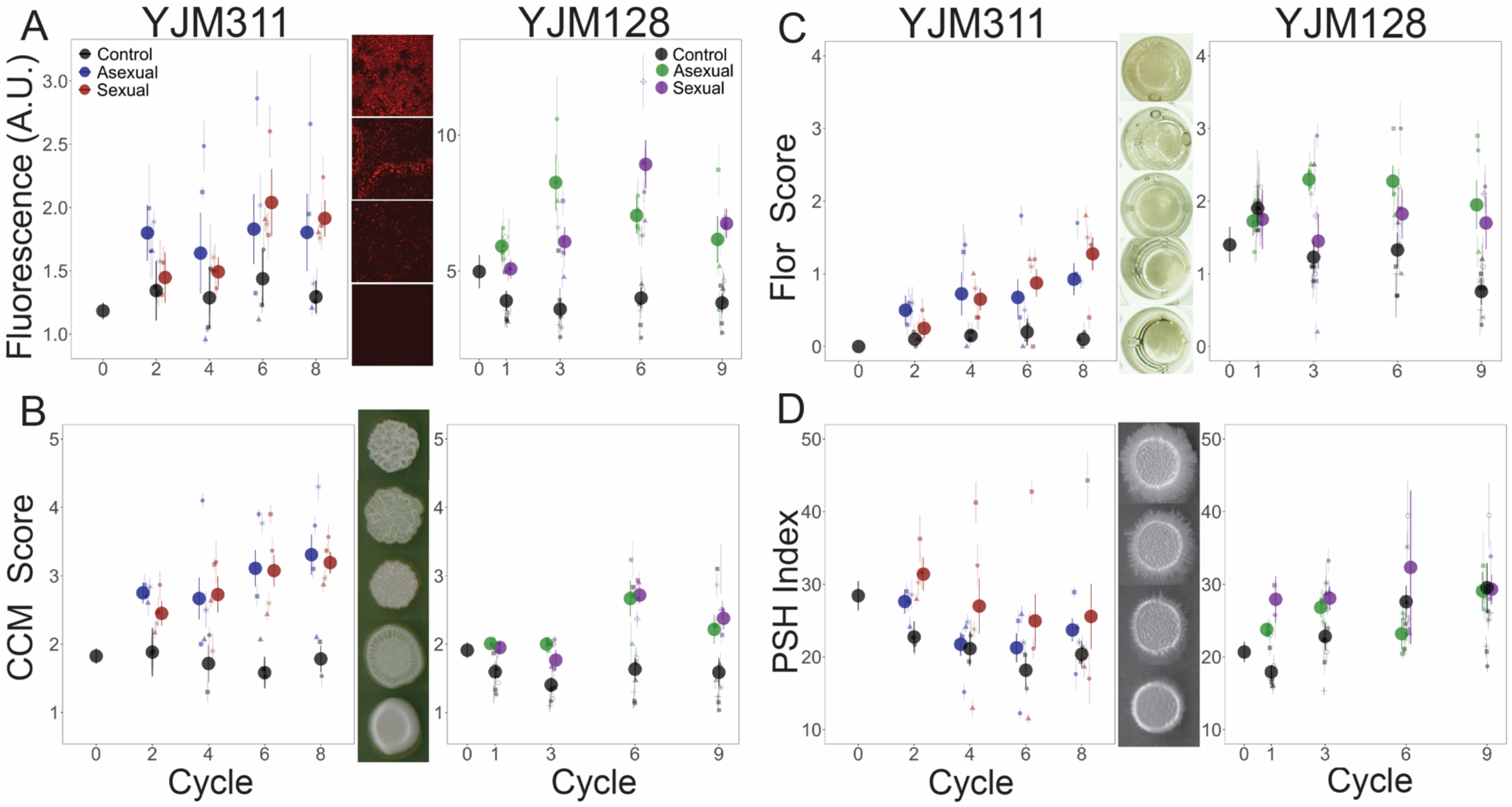
Evolution of Multicellular Phenotypes. Ten clones were isolated from each population at four timepoints and assayed in triplicate (except for flor formation, which had a single replicate). In all panels, large points represent the average of a treatment (asexual, sexual, control) ± 2 s.e.m.; smaller points represent the average of a replicate population ± s.e.m. Data at cycle 0 represent the average of 20 ancestral segregants. Representative images demonstrate the variation found in the phenotypes. **(A)** Plastic adherence was measured using the fluorescence signal of cells remaining in a black, clear-bottom 96-well plate after saturated growth and gentle washing. **(B)** CCM was scored after growth on solid, glucose-limiting medium using the scale on the right, with 1 representing no biofilm and 5 the most structured colonies. **(C)** Flor formation was scored after growth in minimal medium using the scale on the right, with 0 representing no floating cells and 4 representing a full mat. **(D)** PSH was scored after growth on solid nitrogen-limiting medium. Images were processed to determine the percentage of growth pixels that were pseudohyphal compared to the central colony. The trajectory of replicate populations from each ancestral background can be found in Figures S2 and S3.

We next measured the ability of the clonal panel to express three other seemingly different multicellular phenotypes.

### Biofilm Colony Formation

The first multicellular phenotype was the ability to form complex colony morphology (CCM) on solid agar, which is indicative of the ability of a strain to form a differentiated biofilm colony (also known as a “fluffy colony”) [43–47]. This phenotype is correlated with another multicellular phenotype [30], mat formation, which is a biofilm that forms on semi-solid agar [48]; we therefore only assayed CCM. Morphology was scored after growth on solid, glucose-limiting medium using a scale from 1-5, with 1 representing no biofilm and 5 representing the most structured colonies [30,49] (Figure 2B).

In both genetic backgrounds, the selected populations increased in their ability to express CCM compared to the ancestor (Figure 2B), while the control populations either maintained or decreased their expression (YJM311: control*cycle coefficient = -0.024 (confidence interval ±0.056), asexual*cycle = 0.119 (±0.039), sexual*cycle = 0.136 (±0.039); YJM128: control*cycle= 0.007 (±0.020), asexual*cycle = 0.048 (±0.026); sexual*cycle = 0.052 (±0.030); Table S5). YJM311 evolved to express stronger CCM than YJM128, despite the latter evolving for one more cycle.

### Flor Formation

The second multicellular phenotype was the ability to form a flor (or velum), which is a floating mat containing cells attached to one another in an extracellular matrix [29]. Flors form at the liquid-air interface in static conditions and are most commonly found during sherry and wine making processes [50]. Flor formation was scored after growth in minimal medium, with 0 representing no floating cells and 4 representing a full mat. The ability to form flors increased in both genetic backgrounds, despite the cultures being grown with agitation (Figure 2C). In YJM311, the ancestral clones showed no ability to generate flors, yet, its evolved populations did (control*cycle coefficient = 0.007 (confidence interval ±0.048), asexual*cycle = 0.075 (±0.034), sexual*cycle = 0.163 (±0.034); Table S6). In YJM128, ancestral clones showed limited ability to form flors. Its experimental populations either remained or increased in flor-forming ability, while control populations decreased in theirs (control*cycle= -0.109 (±0.032), asexual*cycle = 0.031 (±0.043); sexual*cycle = -0.003 (±0.048); Table S6).

### Pseudohyphal Growth

The final phenotype, pseudohyphal growth (PSH), is a form of filamentous growth thought to represent a foraging strategy. It is characterized by substrate invasion and incomplete separation of mother-daughter cells growing in an elongated, unipolar budding pattern [51]. This phenotype is sometimes correlated with invasive growth, so only PSH was assayed. Filamentous and invasive growth have been associated with pathogenicity and virulence in *S. cerevisiae* [23,24,26], as well as in other fungal pathogens of humans and plants [52]. PSH was scored on solid nitrogen-limiting medium; images were processed to determine the percentage of growth that was pseudohyphal compared to the central colony. Unlike the previously assayed phenotypes, the two genetic backgrounds did not evolve similarly with respect to PSH.

In YJM311, the experimental populations did not increase in their PSH ability compared to the ancestor (Figure 2D); rather, all treatments showed some loss. Throughout the cycles, a moderate level of PSH was maintained in some of the experimental populations, with one sexual replicate doubling its PSH index (Figure S2), but it was lost in the controls and the other experimental populations (control*cycle coefficient = -0.61 (confidence interval ±0.60), asexual*cycle = -0.65 (±0.42), sexual*cycle = -0.97 (±0.43); Table S7).

In YJM128, both the experimental and control populations increased in their PSH ability (control*cycle coefficient = 1.29 (confidence interval ±0.39), asexual*cycle = 0.61 (±0.51), sexual*cycle = 0.91 (±0.55); Table S7). This was true for the control population that was evolved in concert with all experimental populations, as well as the 6 control populations that were initiated subsequently.

The results from both ancestral genetic backgrounds suggest that the evolution of PSH was a response to selection in nutrient limiting conditions and not a response to selection for adherence, as the controls and experimental populations behaved similarly. Of all the adherence and multicellular phenotypes investigated, most of which appeared to increase throughout the experiment, PSH appeared to be independent from the others.

Overall, our phenotyping data show that selection on the ability to adhere to a plastic surface generated a correlated response in multiple multicellular phenotypes, and nutrient limiting conditions favored a further multicellular phenotype in one of the backgrounds.

### Hyper-multicellularity

To understand the phenotypic landscape of the evolved populations and to determine whether the different forms of multicellularity evolved in concert in individual clones, the clonal phenotype data were combined in a principal components analysis (PCA) (Figure 3A, S4, S5). In YJM311, the loadings of the first two components, which explain 78% of the variation, show that evolved clones with the most extreme values of plastic adherence and flor formation do not tend to also excel at PSH. There were clones, however, that evolved to excel in all of the phenotypes, while not obtaining the most extreme values of the individual traits. In YJM128, the first two loadings explain 70% of the variation, and again, PSH appeared separated from the other multicellular phenotypes. Individual correlations between traits bear out this interpretation (Figure S6, S7). When grouped by experimental treatments, clones from control, asexual, and sexual populations tended to occupy their own, somewhat overlapping, phenotypic space (Figures S4 and S5).

**Figure 3:**
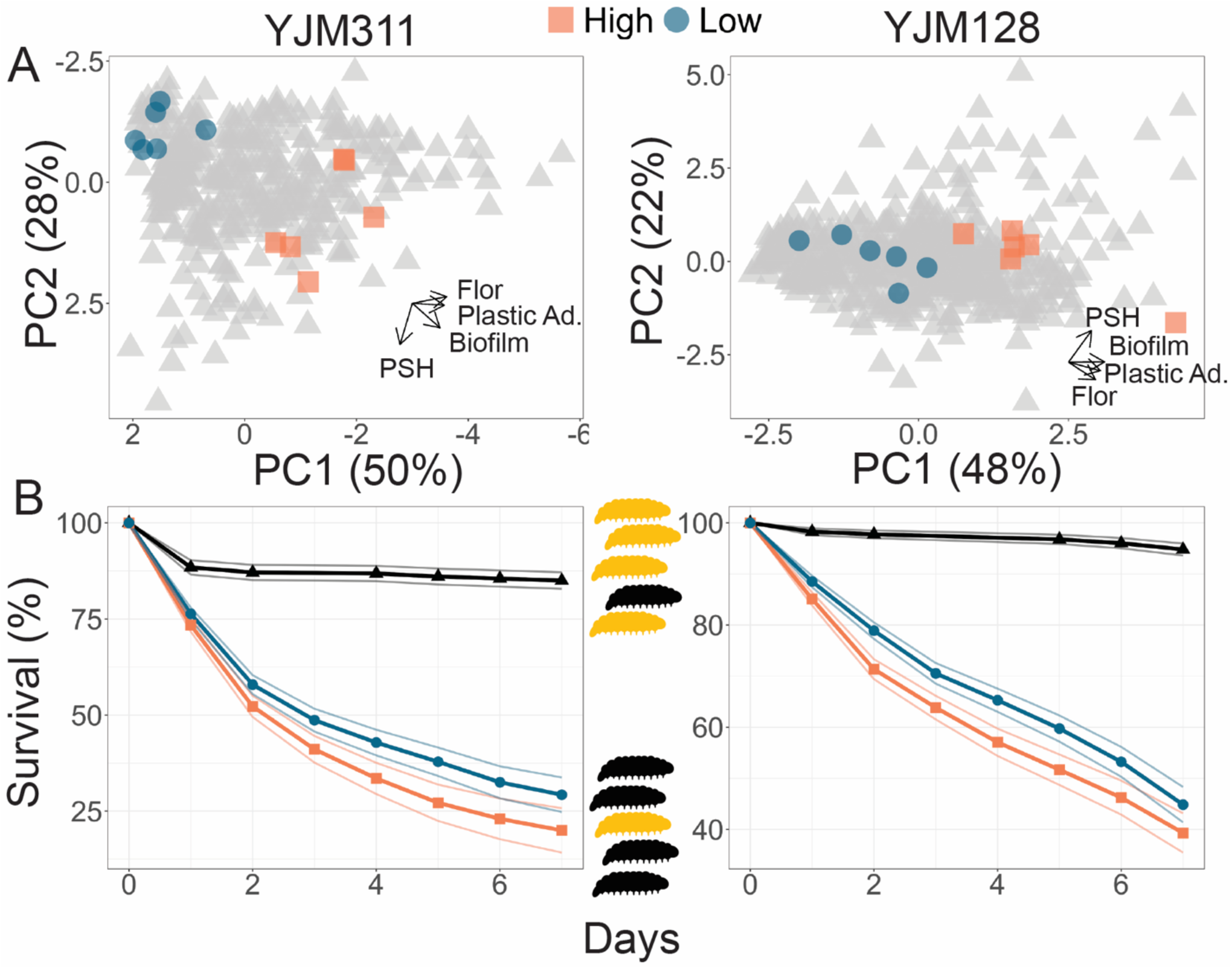
Evolved multicellularity and virulence. **(A)** Principal components analysis of clones from ancestral and evolved populations. Highlighted points represent strains chosen for virulence assays: blue circles represent low multicellularity clones; orange squares represent hyper-multicellular clones; gray triangles represent the rest of the clonal panel. In YJM311, the non-multicellular clones were chosen from ancestral and control populations, while in YJM128, they were chosen from ancestral and early experimental populations. The loadings of PC1 for YJM311 were -0.616\**Flor* - 0.573\**PA* - 0.493\**CCM* + 0.221\**PSH*; for PC2, they were 0.851\**PSH* + 0.506\**CCM* - 0.137\**Flor*. In YJM128, the loadings of PC1 were 0.601\**CCM* + 0.535\**PA* + 0.451\**Flor* + 0.386\**PSH*; for PC2, they were 0.852\**PSH* - 0.226\**PA* - 0.472\**Flor*. PCA with population and cycle information can be found in Figures S4 and S5. **(B)** *G. mellonella* survival curves for strains highlighted in panel A; each strain was injected into 200 larvae for YJM311 derived clones or 180 larvae for YJM128 derived clones. Points represent Kaplan-Meier estimates with confidence limits for non- and hyper-multicellular treatments; black triangles represents the control treatment injected with sterile water. Survival for individual strains is plotted in Figure S10.

In both backgrounds, as the populations evolved, there were individual clones that increased in all abilities, and became “hyper-multicellular”. Thus, a simple process of selection for plastic adherence led to correlated effects in multiple multicellular traits. These correlated effects were apparent both at the population-level, with mean phenotypes increasing in populations over the generations, but also at the individual-level with the evolution of hyper-multicellularity.

### FLO11 *Length Variation*

One possible explanation for the increase in multiple forms of multicellularity is a change in a genetic element common to all four phenotypes. A genome-wide investigation into the genetic basis of three multicellular phenotypes (biofilm formation, PSH, and invasive growth) in a lab strain found that each phenotype appeared to have its own set of hundreds of genes underlying its expression, but also some overlap in select transcription factors and signaling pathways [53]. Notably, the one element that all of the traits had in common, as do other aggregative phenotypes, is the requirement of the cell adhesin, Flo11p [48,54,55], which allows yeast cells to adhere to surfaces and other cells [56].

Flo11p is a cell surface protein with three domains: a C-terminal that facilitates attachment to the cell wall, an exposed N-terminal immunoglobulin-like domain that mediates cell adhesion [57], and a low-complexity, serine-threonine rich B-domain of variable length that extends the adhesion domain away from the cell [56]. The tandem repeats in the B-domain have been shown to be unstable [58,59] and to vary in length naturally [29,60–62]. Differences in the length of this repetitive region have been shown to affect the strength of multicellular phenotypes in some genetic backgrounds [29,59].

To determine whether *FLO11* length changed throughout the experiment, amplicons of the gene were analyzed with electrophoresis in a subset of clones from the final timepoint (Figure 4). In the YJM311 populations, five out of eight experimental populations ended with an approximate 1000bp length increase in some or all clones, while none of the control clones showed an increase in length. It is unknown whether the change in length was due to independent *de novo* mutations or selection favoring an existing allele. The similar allelic length in multiple replicate populations favors the latter explanation. It is possible that during the generation of the starting recombinant pool, there was a mutation that was not detected in the subset of ancestral clones later chosen for analysis. In this genetic background, *FLO11* length is not correlated with the strength of plastic adherence, nor with the other three multicellular phenotypes (Figure S8).

**Figure 4:**
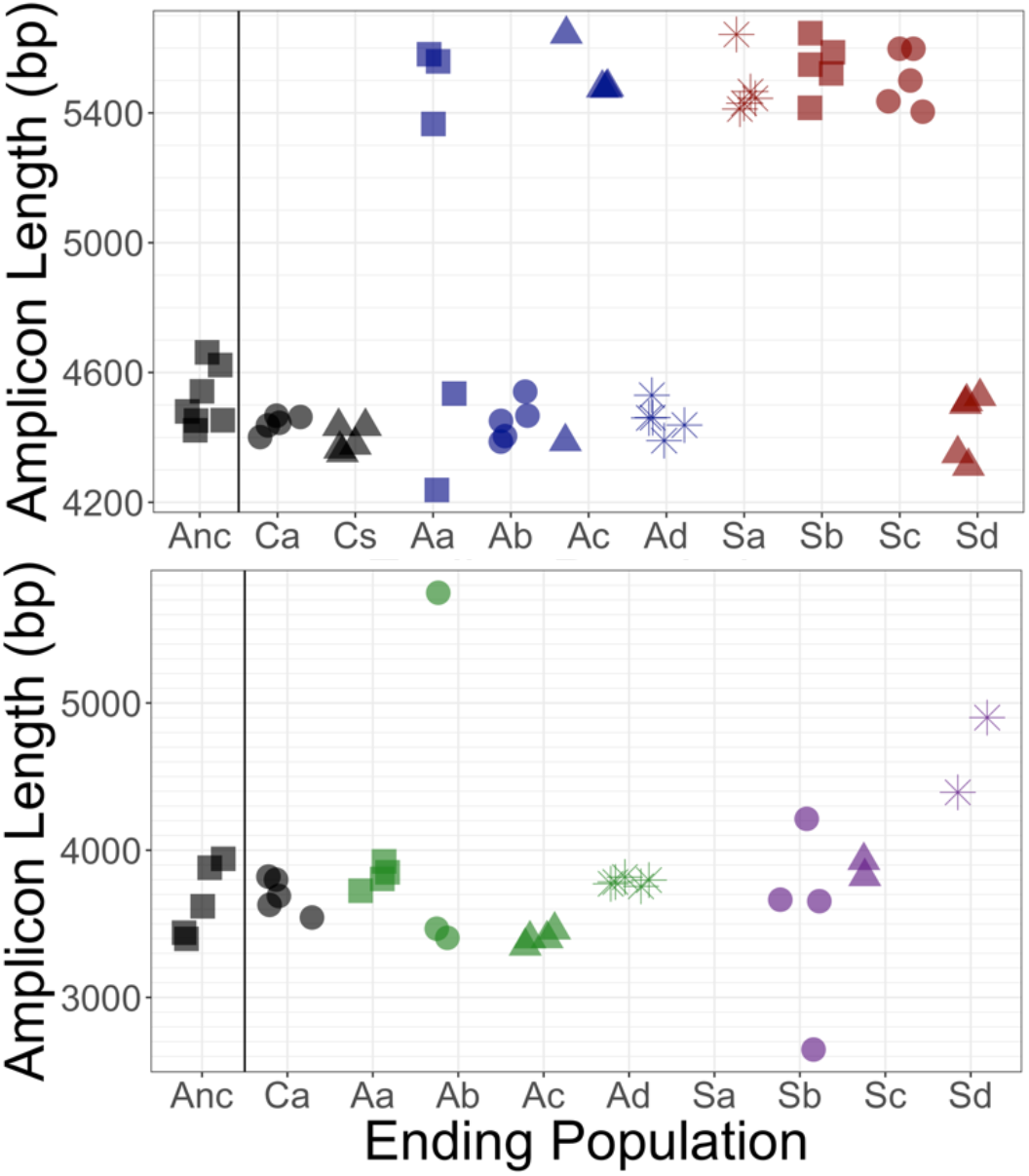
*FLO11* length evolution. In 8 ancestral clones and 5 clones per replicate population at the final cycle, the full gene was amplified and run through a BioAnalyzer to determine its length. Amplicons of this length have an accuracy of ±100 bp. x-axis: Anc refers to ancestor, C refers to control populations, A to asexual populations, and S to sexual populations; a-d denote replicates. Top panel: In YJM311, it appears there were two major length alleles, with the possibility of derived variants with smaller changes in length. Bottom panel: In YJM128, it appears there were also two alleles, separated by ∼500 bp. Clones from the final timepoint show variation in length.

In YJM128, the ancestral pool likely had two alleles separated by a few hundred basepairs. The ending asexual populations appeared to have these alleles, with one much longer allele in a clone in one replicate. The ending sexual populations contained the ancestral alleles, as well as other variants both longer and shorter. Clones from one replicate could not be amplified (Sa), suggesting the possibility of a mutation in the region where the primers anneal. Again, *FLO11* length was not correlated with the strength of plastic adherence, nor with the other three multicellular phenotypes (Figure S9).

Thus, while *FLO11* length evolved during the experiment, it does not appear to be the cause of the correlated response to selection on adherence. However, this does not rule out the possibility that *FLO11* plays a role. It is possible that expression of the gene, through its complex regulatory network [63–65], is related to the phenotypic response to adherence selection.

### Virulence

We sought to determine if the evolved changes had an effect on virulence. Virulence was measured using larvae of the greater wax moth, *Galleria mellonella*, an invertebrate model used to study microbial pathogenesis and virulence [66], including in *S. cerevisiae* [67]. Using the PCA results as a guide, for each genetic background, we identified six hyper-multicellular clones from later time points, and six non-multicellular clones from throughout the experiment (Figure 3A, Tables S1, S2). Strains were grown in the medium in which they were evolved, then washed, adjusted for density, and injected into larvae. Larval survival and pupation were monitored for the next 7 days. Different batches of larvae can be variable in their response to microbial insult; therefore, to ensure reproducibility of results, the experiment was repeated multiple times with numerous batches of larvae, with each batch being challenged by all strains from a genetic background. For the clones derived from YM311, of the 2400 larvae injected with yeast, 1809 did not survive through day 7. Larvae injected with a hyper-multicellular strain were 1.28 times more likely to die than those injected with a non-multicellular strain (*p* < 0.001) (Figures 3B, S10). For the clones derived from YM128, of the 2160 larvae injected with yeast, 1113 did not survive through day 7. Larvae injected with a hyper-multicellular strain were 1.29 times more likely to die than those injected with a non-multicellular strain (*p =* 0.032) (Figures 4B, S6).

Thus, the evolutionary changes brought about by selection for adherence to a plastic bead, led to the incidental evolution of increased virulence.

## Discussion

Our results demonstrate that selection on one yeast trait can generate a correlated response in other traits— a common feature of organismal evolution [68]—, but here, the correlated traits may have included those associated with virulence. In this experiment, favoring the ability to adhere to plastic, a surface that is alarmingly common in industrial, medical, and domestic settings [69], led to a suite of aggregative phenotypes and increased virulence.

The accidental virulence hypothesis proposes that selection for survival in harsh conditions may lead to traits that predispose microbes to virulence. However, harsh environmental conditions can also favor traits that favor the collective, or multicellular phenotypes [70], so it is perhaps not surprising that other forms of multicellularity increased throughout this experiment. Furthermore, in its long evolutionary history, *S. cerevisiae* has evolved the genetic capability to express multiple different multicellular phenotypes, most of which are induced in nutrient limiting conditions; the experiment presented here was performed in such conditions (glucose limited medium). Other experiments with yeast growing in nutrient limiting conditions have also resulted in the unintended evolution of aggregative behaviors [71]. In this light, the correlated phenotypic responses in this experiment are not entirely unexpected. However, the evolution of *multiple* multicellular phenotypes in *two* independent genetic backgrounds was not anticipated.

Previous research on a panel of environmental isolates found no correlation between the phenotypes assayed here [30], and in a tractable lab strain capable of aggregative behaviors, each phenotype was associated with its own set of genes [53]. However, there is overlap in the requirement of *FLO11* and its regulators, and despite the fact that the phenotypes are induced by different nutrient signals, there are numerous conserved signaling pathways contributing to filamentous, multicellular growth of all forms (e.g., cAMP-PKA, TOR, filamentous MAPK, Rim101) [31,72]. It is well known that genetic background and genetic architecture can have strong effects on the expression and correlation of traits [73]. In the case of the filamentous phenotypes assayed here, a genetic background that contains variants in the main signaling pathways may lead to a correlation of the phenotypes, while variants expressed later in the development of the phenotype, that are specific to a single trait, may not lead to such a correlation. The effect of the different types of genetic variants suggests that some strains and genetic backgrounds are more likely to evolve virulence from selection in the open environment.

The strains used in this experiment each contain ∼50,000 heterozygous sites and differ from each other by ∼25,000 SNPs. It is possible that they contained genetic variation in canonical signaling pathways, allowing for the evolution of hyper-multicellular strains. Interestingly, in both backgrounds, pseudohyphal growth appeared to evolve independently of the other phenotypes. Future research will investigate the sorting of the genetic variation, as well as the new mutations, that led to the observed phenotypic evolution in these populations.

In our experiment, it is unclear which trait was associated with increased virulence: plastic adherence, a different multicellular trait, general hyper-multicellularity, or an entirely different trait unintentionally favored in the experiment. Regardless of the specific trait causing increased virulence, the experiment demonstrates that selection for a specific trait in an environment that is entirely devoid of host organisms can still inadvertently lead to virulence and pathogenicity.

The role of environmental opportunistic pathogens in infectious disease is important to consider as humans generate novel ecological niches through the use of plastics, encroach on more habitats, and especially, as the global climate changes [7]. Warmer water temperatures and climate disruptions have been linked to the incidence of illness caused by the marine bacterium *Vibrio vulnificus* [74], and an increase in global temperature has been hypothesized to be related to the simultaneous emergence of *C. auris* infections on multiple continents [75,76]. More generally, the narrowing of the gap between mammalian body temperatures and the ambient environment may create opportunities for fungi to exploit new host niches [6]. Thus, as new selective pressures act on existing abundant genetic variation, there is the opportunity to create unintended, accidental pathogens.

## Materials and Methods

### Strains

To generate strains appropriate for downstream phenotyping assays, the original diploid isolates were was engineered to express a fluorescence protein by fusing mCherry to the C-terminal region of the highly expressed *PGK1* gene, generating HMY7 (YJM311 *PGK1-mCherry-KanMX*) [77] and HMY355 (YJM128 *PGK1-mCherry-HygMX*). After being subject to selection for 8-9 cycles, clones with different multicellular phenotypes were isolated from each replicate population (Tables S1-S2).

### Media

Experimental populations were grown in Evolution Medium (EM; 0.17% yeast nitrogen base without ammonium sulfate and without amino acids, 0.1% glutamic acid, 0.1% dextrose) supplemented with G418 (200 μg/ml) or Hygromycin B (300 μg/ml). Cells were sporulated on solid medium (1% potassium acetate, 2% agar) and digested using an overnight zymolyase-β-glucuronidase procedure [38,78]. Phenotypes were assayed on YPD (1% yeast extract, 2% peptone, 2% dextrose, 2% agar), low dextrose (LD) YPD (0.1% dextrose), 2X SLAD (0.34% yeast nitrogen base without ammonium sulfate and without amino acids, 2% dextrose, 50 μmol ammonium sulfate, 2% agar), or in liquid SD (0.17% yeast nitrogen base without amino acids and with ammonium sulfate, 2% dextrose).

### Experimental Evolution

HMY7 and HMY355 were grown in 10ml YPD, sporulated, digested, grown to saturation in 10ml EM, and used to inoculate 10 replicate populations: 4 sexual, 4 asexual, and 1 control of each reproductive type.

Experimental populations derived from YJM311 were evolved for 8 12-day cycles, for a total of ∼350 generations; populations from YJM128 were evolved for 9 cycles, for a total of ∼400 generations. In each cycle, populations were grown in 10ml of EM in a glass tube containing a sterile 7mm polystyrene bead (American Education Products), population size ∼ 2 × 10^8^. After 48 h at 30°C in a rotator drum, the bead was removed with sterile disposable forceps, washed twice, suspended in 500ul of sterile H_2_O in a microcentrifuge tube, and gently sonicated (UP200St with VialTweeter, Heischler Ultrasound Technology) to detach cells from the bead. The cell suspension was used to inoculate the next 10ml EM tube. The number of cells on the bead varied over the experiment (Figure 1B). After 4 serial transfers, asexual populations were refrigerated and sexual populations were sporulated for 48 h. Asci were digested overnight, and the spores resuspended in 1ml of EM to allow germination and mating (population size ∼ 10^5^ spores). Finally, the refrigerated cultures and the mated spores were used to begin the next 12-day cycle (Figure S1).

### Population Phenotyping

To estimate adherence evolution, all populations from all cycles were assayed using the same batch of medium. 10ml EM cultures were inoculated with cryopreserved glycerol stocks and grown for 48 h. From these, two replicate test tubes were inoculated with two beads in each, for a total of four beads per population per time point. The cultures were grown and the beads processed as in the experimental cycle; cell counts were made using a hemocytometer with the sonicated cell suspension. This entire process was repeated a second time, for a total of 8 beads per population per cycle for YJM311 populations.

### Clonal Phenotyping

Twenty clones were isolated from the ancestral population and 10 clones were isolated from each replicate population at four cycle timepoints: 2, 4, 6, 8, for YJM311, and 1, 3, 6, 9, for YJM128. The clonal strains were arrayed in a 96-well format and cryopreserved. To assay social phenotypes, saturated YPD cultures were resuspended and pinned to different media using a 96-pin multi-blot replicator (V&P Scientific no. VP408FP6), wrapped in parafilm, and incubated at 30°C.

### Plastic Adherence

Clones were grown in 200μl EM for 48h in 3 replicate black, clear-bottom, non-treated 96-well plates. Optical density was measured, then culture was removed, and plates were gently washed with water three times and dried upside down for 1h. Fluorescence readings were taken with a Spectramax M2e (Molecular Devices) and used as a proxy for the number of cells that remained attached to the wells. To account for differences in growth, each fluorescence reading was divided by the optical density of the well.

### Flor formation

Clones were grown in 200μl SD for 5d and imaged on an Olympus SZX16 dissecting scope. Flor formation was scored using the scale in Figure 2.

### Complex colony morphology (CCM)

Clones were pinned to 3 replicate LD omni trays, incubated for 7 days, and imaged on an EPSON Expression 11000 XL scanner. Colonies were scored for complexity using the scale in Figure 2.

### Pseudohyphal growth

Clones were pinned to 3 replicate 2X SLAD omni trays, incubated for 8 days, and scanned. Images were processed using a custom script that determined the percentage of colony pixels comprising the pseudohyphae [35].

### Data Analysis

Bead cell count data from experimental evolution were log-transformed and analyzed using a mixed effects linear model in R [79] with the lme4 package [80]. Replicate populations within treatments (control, asexual, sexual) was considered a random effect. Because all populations were begun from a single ancestral pool, the intercept was set as the mean value of the ancestor and not allowed to vary among treatments. Therefore, the only fixed effect was the interaction between cycle and treatment, which tested the differences among the slopes of the three treatments. Clonal data were analyzed similarly, with the untransformed average score of a phenotype as the independent variable. Finally, the average phenotyping data for each clone were combined for a principal components analysis in R using the *princomp* function. Figures were produced using ggplot2 [81].

### Virulence Assay

10ml EM cultures of evolved strains (Table S1) were grown for 48h, washed and resuspended in sterile water to a concentration of 10^9^ cells/ml based on hemocytometer counts. 4 μl of culture or control water was injected into the final posterior proleg of *Galleria mellonella* larvae (Vanderhorst Wholesale Inc., https://www.waxworms.net) weighing between 9 and 14 mg using a Hamilton PB600-1 Repeating Dispenser with a 27-gauge needle. Each strain was injected into 20 larvae on the same day using the same shipment of *G. mellonella*; 20 control larvae were injected at the start of the assay and at the end. The same assay was repeated the next day with the same shipment of larvae, for a total of 40 larvae/strain/shipment. Multiple shipments were used for the virulence measurements, for a total of 200 larvae per strain for YJM311-derived strains and 160 larvae per strain for YJM128-derived strains. After injection, larvae were incubated at 30°C and survival was monitored for 7 days; larvae that turned black and no longer responded to tactile stimulation were considered dead and removed from the population, as were larvae beginning to pupate.

Data were analyzed with a mixed effects Cox model using the coxme package [82] in R [79]. Death was recorded as the day larvae were removed from the population; larvae were censored if removed for pupation. The model included treatment (high vs. low multicellular) as a fixed effect, and strain and larval batch as random effects.

### FLO11 Length

Of the clones assayed for multicellular phenotypes, 8 ancestral clones and 5 clones from the final time point of each replicate population were chosen for length analysis. Genomic DNA was extracted using the MasterPure Yeast DNA Purification Kit (Lucigen). *FLO11* was amplified with Phusion polymerase (New England BioLabs) and primers targeting the entire gene (forward: GCC TCA AAA ATC CAT ATA CGC ACA CTA TG, reverse: TTA GAA TAC AAC TGG AAG AGC GAG TAG). Cycle conditions followed manufacturers recommendations and included a melting temperature of 58°C and 3-min extension time. Gene length was estimated by running PCR amplicons through the Agilent 2100 BioAnalyzer using the Agilent DNA 7500 kit (as in ref [60]).

## Acknowledgements

We thank Paul Magwene for strains, and Joseph Heitman and Anna Averette for guidance with *G. mellonella* assays. The research was funded by National Institutes of Health grant R15GM122032 and National Science Foundation grant DEB-1839555 to HAM, and William & Mary Charles Center Summer Fellowships to LIE, DM, DVM, and JAS.

---

**Table S1:**
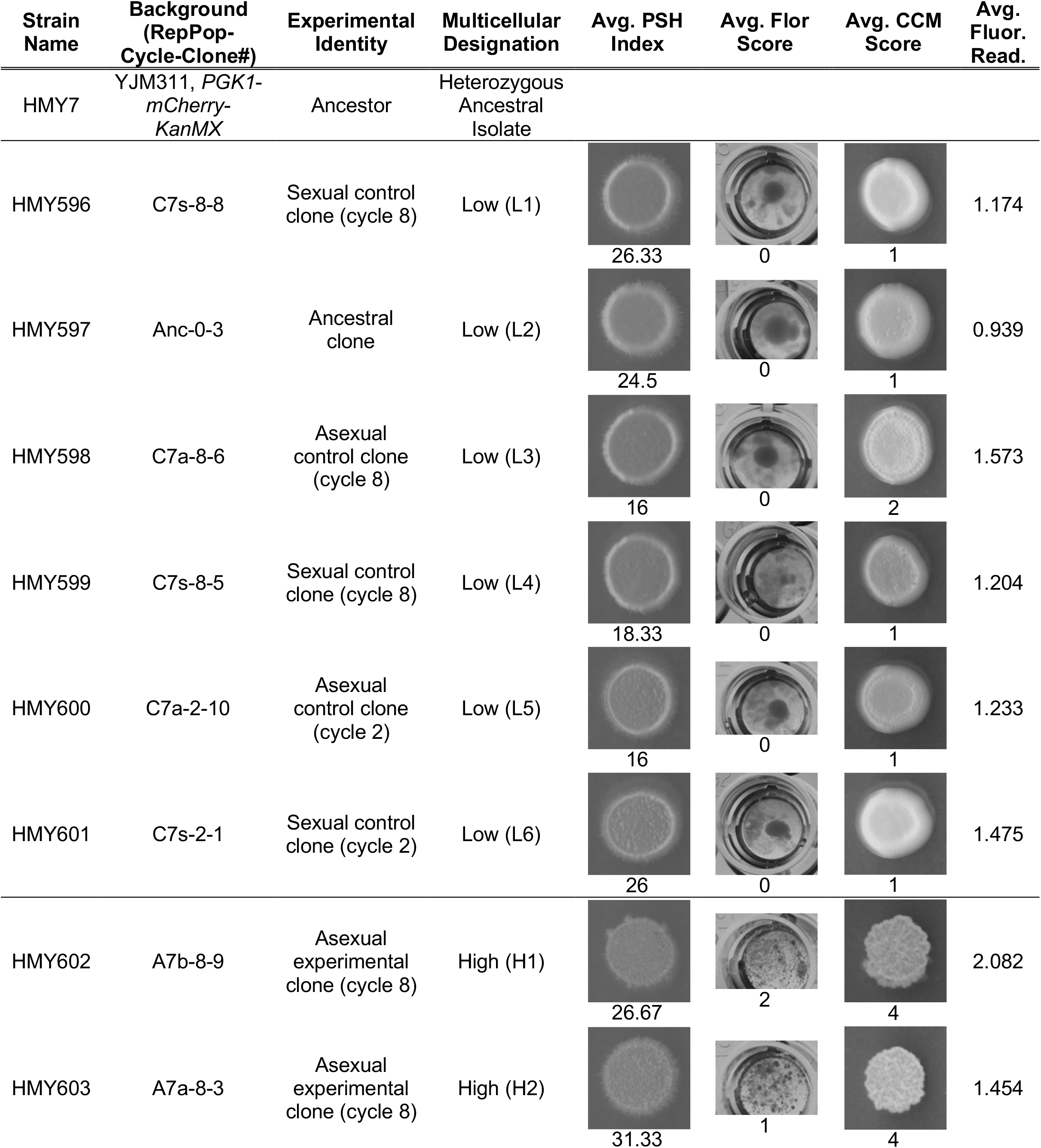
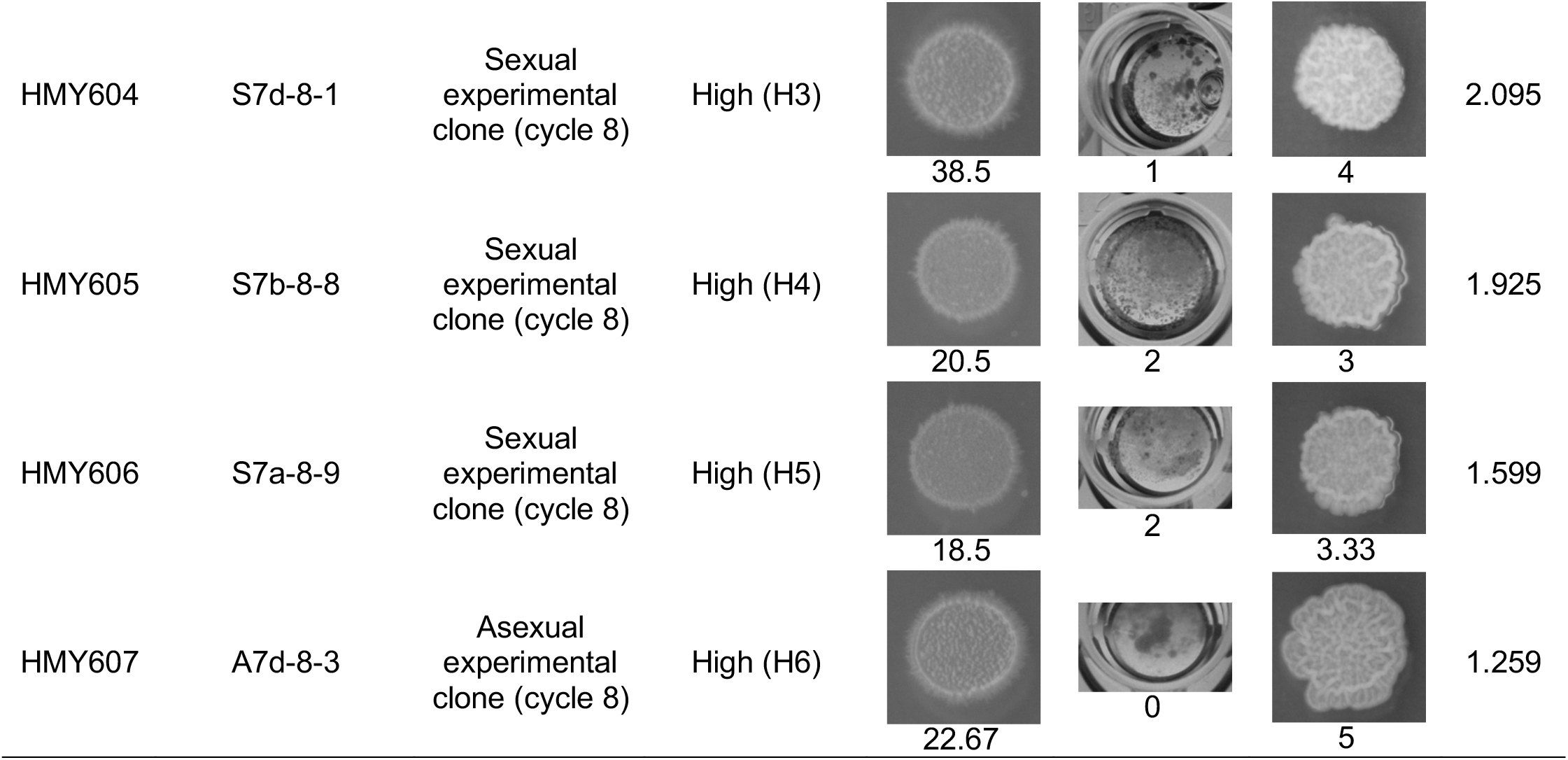
YJM311 Strains used in virulence experiments.

| Strain Name | Background (RepPop-Cycle-Clone#) | Experimental Identity | Multicellular Designation | Avg. PSH Index | Avg. Flor Score | Avg. CCM Score | Avg. Fluor. Read. |
| --- | --- | --- | --- | --- | --- | --- | --- |
| HMY7 | YJM311, *PGK1-mCherry-KanMX* | Ancestor | Heterozygous Ancestral Isolate |  |  |  |  |
| HMY596 | C7s-8-8 | Sexual control clone (cycle 8) | Low (L1) | 26.33 | 0 | 1 | 1.174 |
| HMY597 | Anc-0-3 | Ancestral clone | Low (L2) | 24.5 | 0 | 1 | 0.939 |
| HMY598 | C7a-8-6 | Asexual control clone (cycle 8) | Low (L3) | 16 | 0 | 2 | 1.573 |
| HMY599 | C7s-8-5 | Sexual control clone (cycle 8) | Low (L4) | 18.33 | 0 | 1 | 1.204 |
| HMY600 | C7a-2-10 | Asexual control clone (cycle 2) | Low (L5) | 16 | 0 | 1 | 1.233 |
| HMY601 | C7s-2-1 | Sexual control clone (cycle 2) | Low (L6) | 26 | 0 | 1 | 1.475 |
| HMY602 | A7b-8-9 | Asexual experimental clone (cycle 8) | High (H1) | 26.67 | 2 | 4 | 2.082 |
| HMY603 | A7a-8-3 | Asexual experimental clone (cycle 8) | High (H2) | 31.33 | 1 | 4 | 1.454 |
| HMY604 | S7d-8-1 | Sexual<br>experimental<br>clone (cycle 8) | High (H3) | 38.5 | 1 | 4 | 2.095 |
| HMY605 | S7b-8-8 | Sexual<br>experimental<br>clone (cycle 8) | High (H4) | 20.5 | 2 | 3 | 1.925 |
| HMY606 | S7a-8-9 | Sexual<br>experimental<br>clone (cycle 8) | High (H5) | 18.5 | 2 | 3.33 | 1.599 |
| HMY607 | A7d-8-3 | Asexual<br>experimental<br>clone (cycle 8) | High (H6) | 22.67 | 0 | 5 | 1.259 |

**Table S2:**
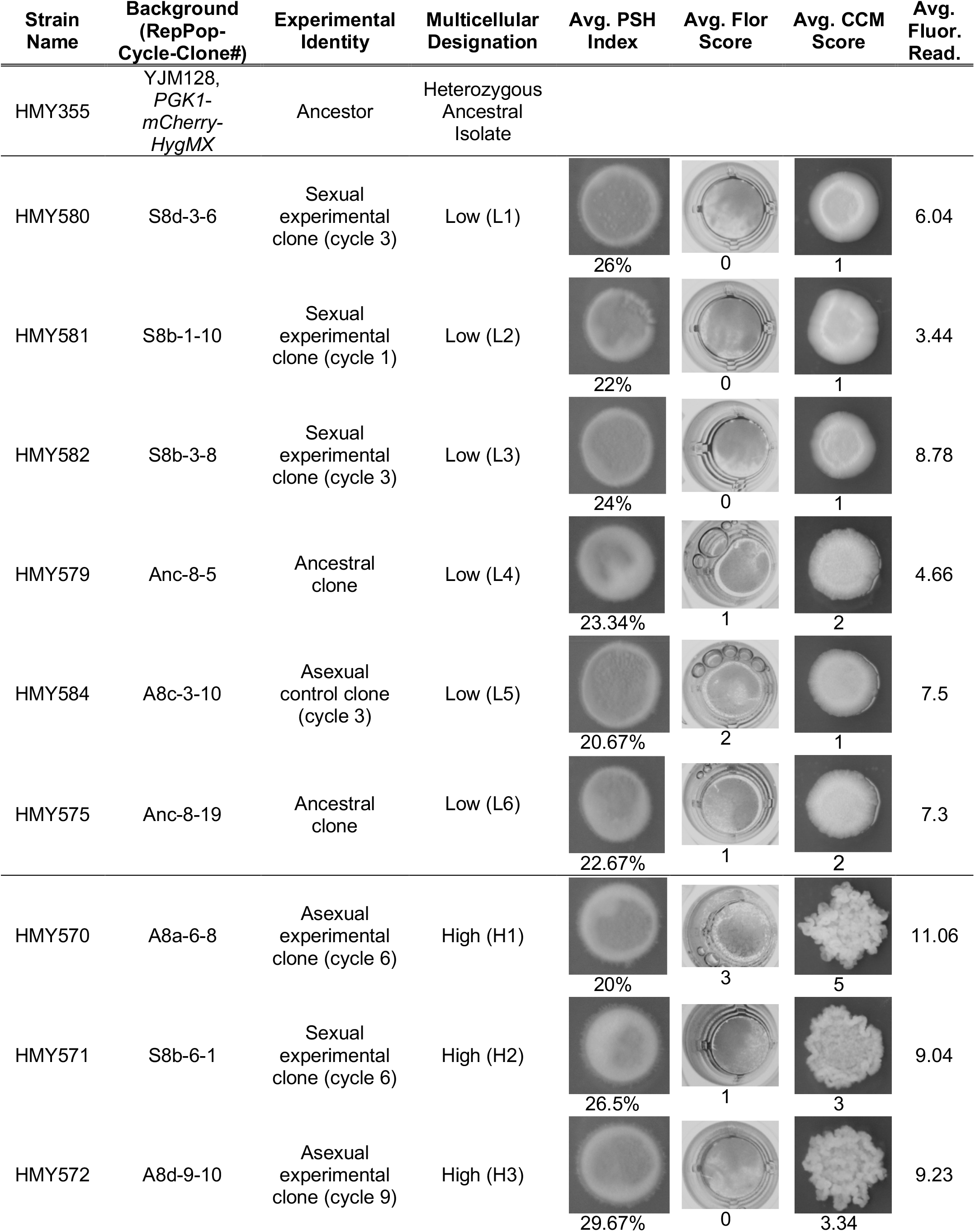
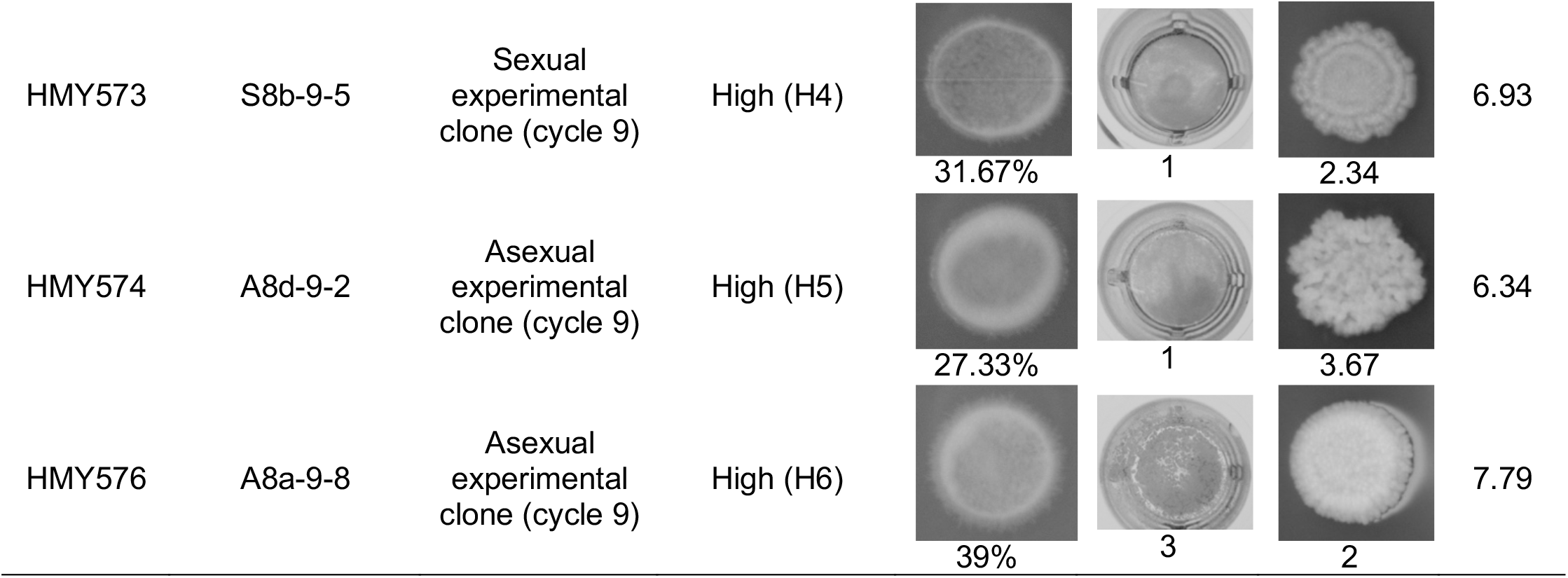
YJM128 Strains used in virulence experiments.

**Table S3:** Results of the mixed-effect linear model for whole-population cell count data over experimental cycles. Cell counts were transformed by adding one and taking the natural log; data were analyzed with the lme4 package in R. Coefficients whose confidence intervals do not encompass zero are bolded.

|  | YJM311 |  |  | YJM128 |  |  |
| --- | --- | --- | --- | --- | --- | --- |
| **<u>Fixed Effects</u>** | <u>coefficient</u> | <u>st. error</u> | <u>t-value</u> | <u>coefficient</u> | <u>st. error</u> | <u>t-value</u> |
| Cycle: Control | -0.05528 | 0.03437 | -1.608 | 0.02182 | 0.09041 | 0.241 |
| Cycle: Sexual | **0.05407** | 0.02433 | 2.223 | **0.46993** | 0.12151 | 3.868 |
| Cycle: Asexual | **0.18435** | 0.02422 | 7.612 | **0.59841** | 0.12747 | 4.695 |
| **<u>Random Effects</u>** | <u>variance</u> |  |  | <u>variance</u> |  |  |
| Population(Treatment) | 0.2593 |  |  | 7.683 |  |  |
| Residual | 0.9466 |  |  | 17.302 |  |  |
| Residual DF | 629 |  |  | 525 |  |  |

**Table S4:** Results of the mixed-effect linear model for clonal plastic adherence data. Data were analyzed with the lme4 package in R. Coefficients whose confidence intervals do not encompass zero are bolded.

|  | YJM311 |  |  | YJM128 |  |  |
| --- | --- | --- | --- | --- | --- | --- |
| **<u>Fixed Effects</u>** | <u>coefficient</u> | <u>st. error</u> | <u>t-value</u> | <u>coefficient</u> | <u>st. error</u> | <u>t-value</u> |
| Cycle: Control | 0.008143 | 0.028509 | 0.286 | 0.01614 | 0.03805 | 0.424 |
| Cycle: Sexual | 0.039812 | 0.020266 | 1.964 | 0.01174 | 0.05184 | 0.227 |
| Cycle: Asexual | **0.100228** | 0.020125 | 4.980 | **0.18686** | 0.05661 | 3.301 |
| **<u>Random Effects</u>** | <u>variance</u> |  |  | <u>variance</u> |  |  |
| Population(Treatment) | 0.1482 |  |  | 2.328 |  |  |
| Residual | 0.4347 |  |  | 4.066 |  |  |
| Residual DF | 391 |  |  | 574 |  |  |

**Table S5:** Results of the mixed-effect linear model for clonal CCM data. Data were analyzed with the lme4 package in R. Coefficients whose confidence intervals do not encompass zero are bolded.

|  | YJM311 |  |  | YJM128 |  |  |
| --- | --- | --- | --- | --- | --- | --- |
| **<u>Fixed Effects</u>** | <u>coefficient</u> | <u>st. error</u> | <u>t-value</u> | <u>coefficient</u> | <u>st. error</u> | <u>t-value</u> |
| Cycle: Control | -0.02432 | 0.02783 | -0.874 | 0.007192 | 0.009940 | 0.724 |
| Cycle: Sexual | **0.11899** | 0.01968 | 6.046 | **0.048360** | 0.013161 | 3.675 |
| Cycle: Asexual | **0.13610** | 0.01968 | 6.916 | **0.052230** | 0.014772 | 3.536 |
| **<u>Random Effects</u>** | <u>variance</u> |  |  | <u>variance</u> |  |  |
| Population(Treatment) | 0.2748 |  |  | 0.1702 |  |  |
| Residual | 0.3588 |  |  | 0.2789 |  |  |
| Residual DF | 395 |  |  | 584 |  |  |

**Table S6:**
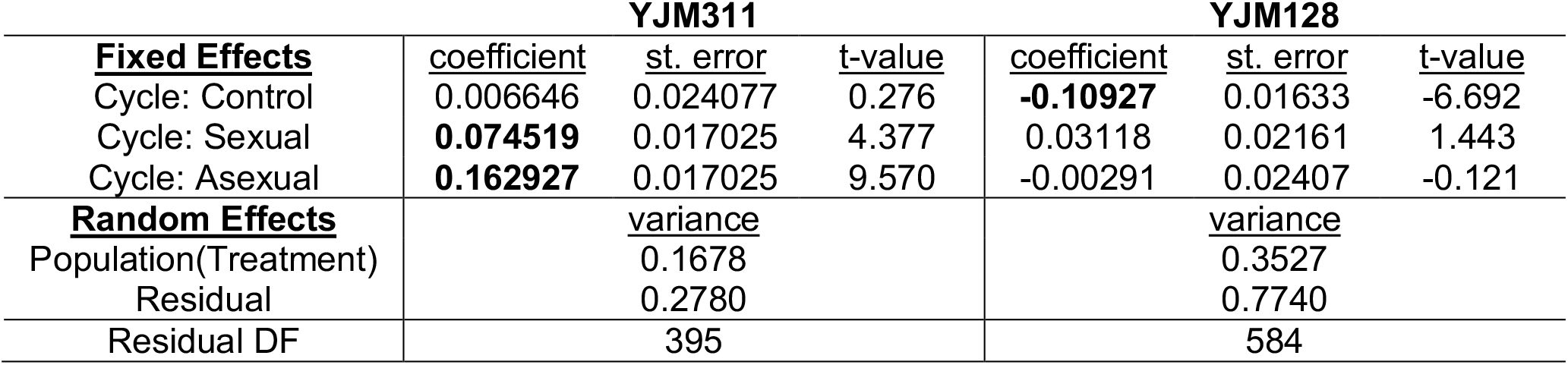
Results of the mixed-effect linear model for clonal flor data. Data were analyzed with the lme4 package in R. Coefficients whose confidence intervals do not encompass zero are bolded.

**Table S7:**
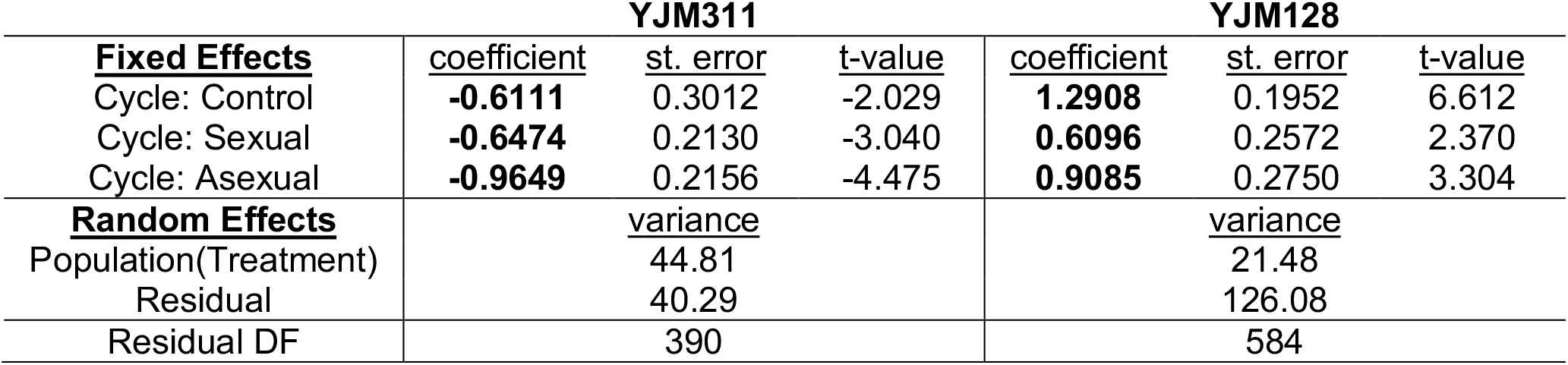
Results of the mixed-effect linear model for clonal PSH data. Data were analyzed with the lme4 package in R. Coefficients whose confidence intervals do not encompass zero are bolded.

**Figure S1:**
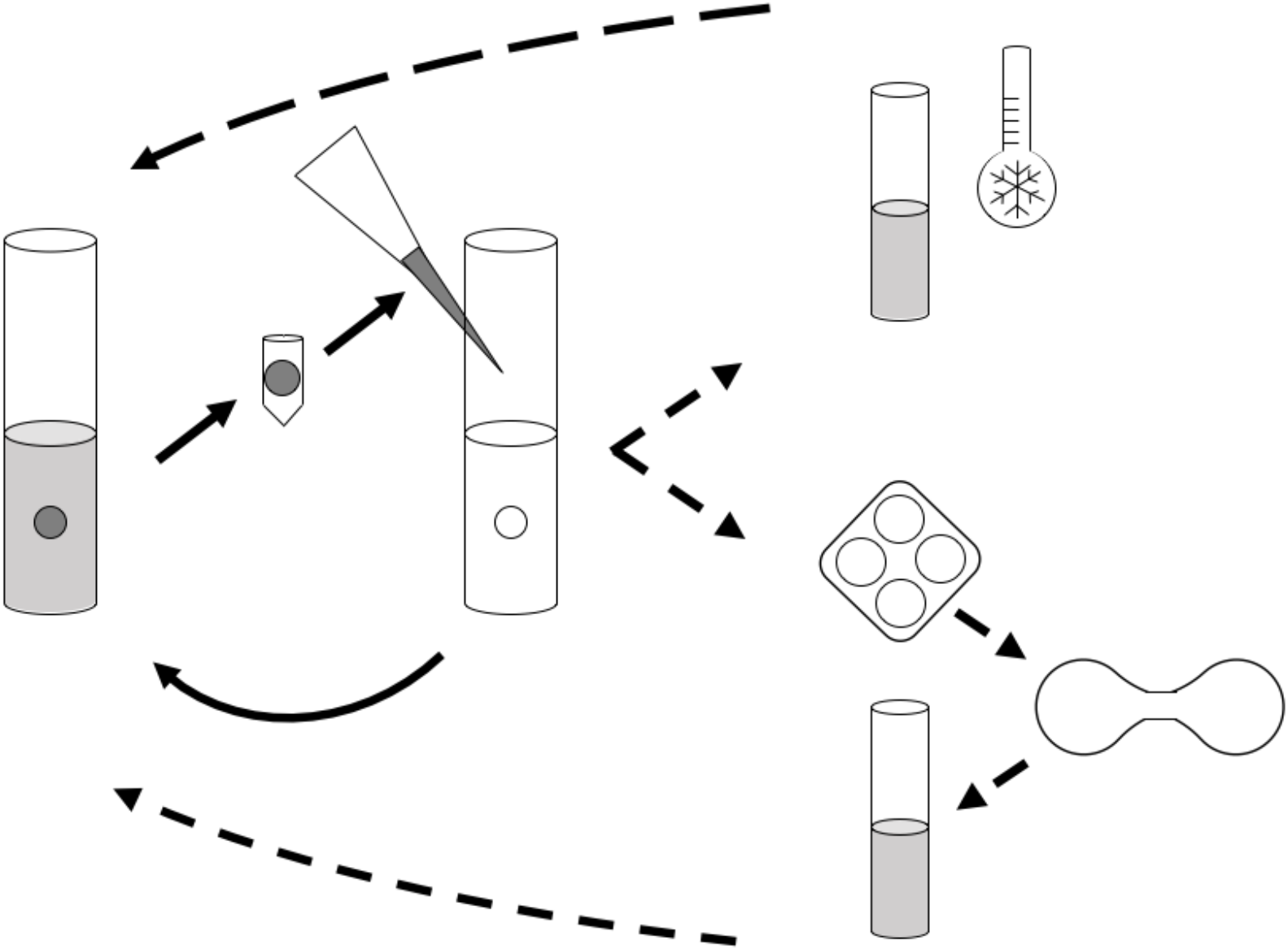
Schematic of the Experimental Cycle. Solid arrows represent steps in serial transfer; dashed lines represent punctuated sexual treatments. Populations were evolved on a 12-day cycle. In each cycle, populations were grown in 10ml of minimal medium in a glass tube containing a sterile 7mm polystyrene bead for 48 h with rotation. The bead was removed, washed, suspended in water, and gently sonicated to detach cells from the bead. The cell suspension was used to inoculate the next 10ml tube. After 4 serial transfers, asexual populations were refrigerated and sexual populations were sporulated for 48 h. Asci were digested, germinated and mated. Refrigerated cultures and mated spores were used to begin the next 12-day cycle.

**Figure S2:**
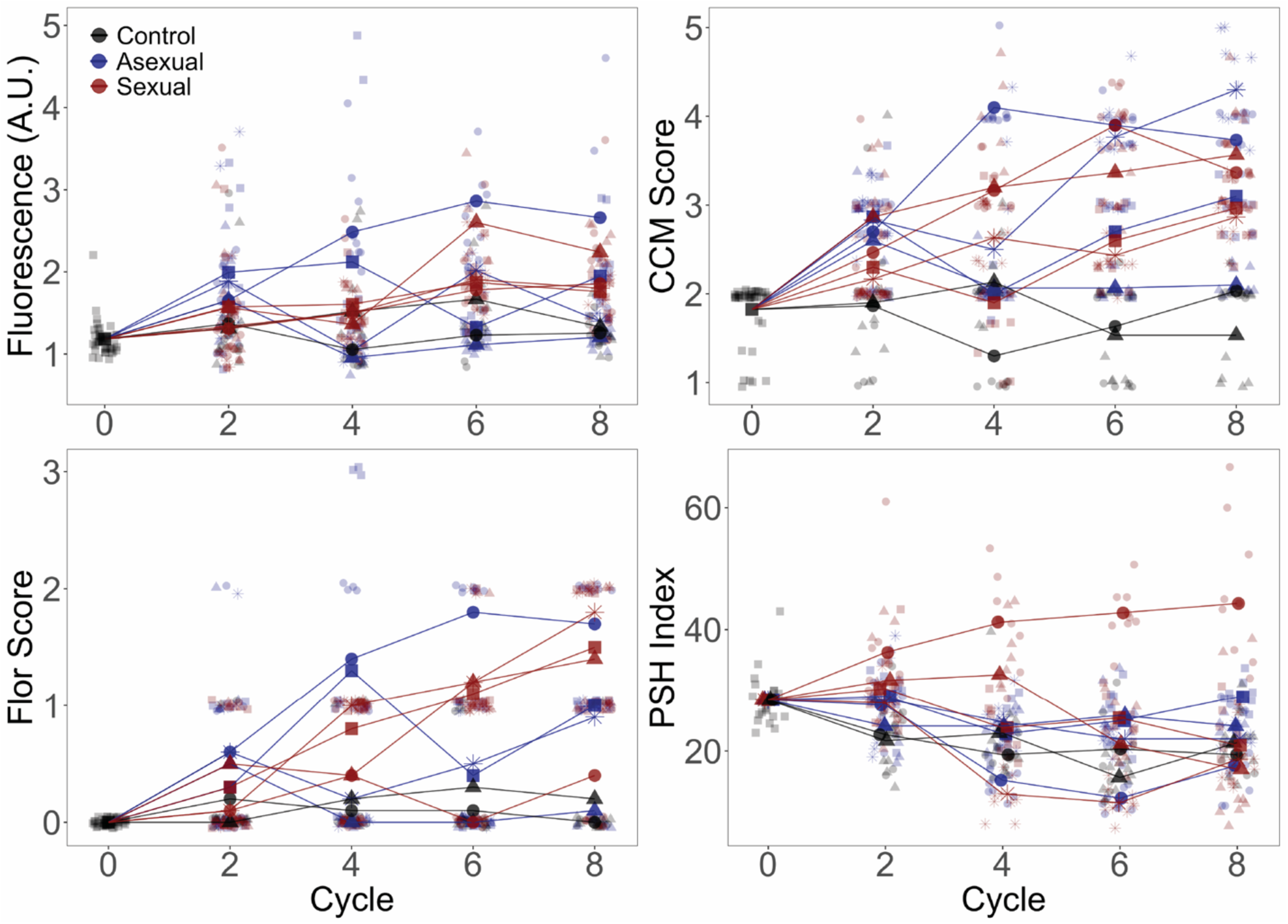
Evolved Phenotypes by Replicate Population for the YJM311 Background. Phenotypes were assessed as in Figure 2. Larger points represent replicate population averages, with each replicate represented by a different shape/color combination (as in Figure 4); smaller points represent measurements from individual clones. Fluorescence, CCM, and PSH are average values of three replicate measurements per clone; Flor is based on one replicate.

**Figure S3:**
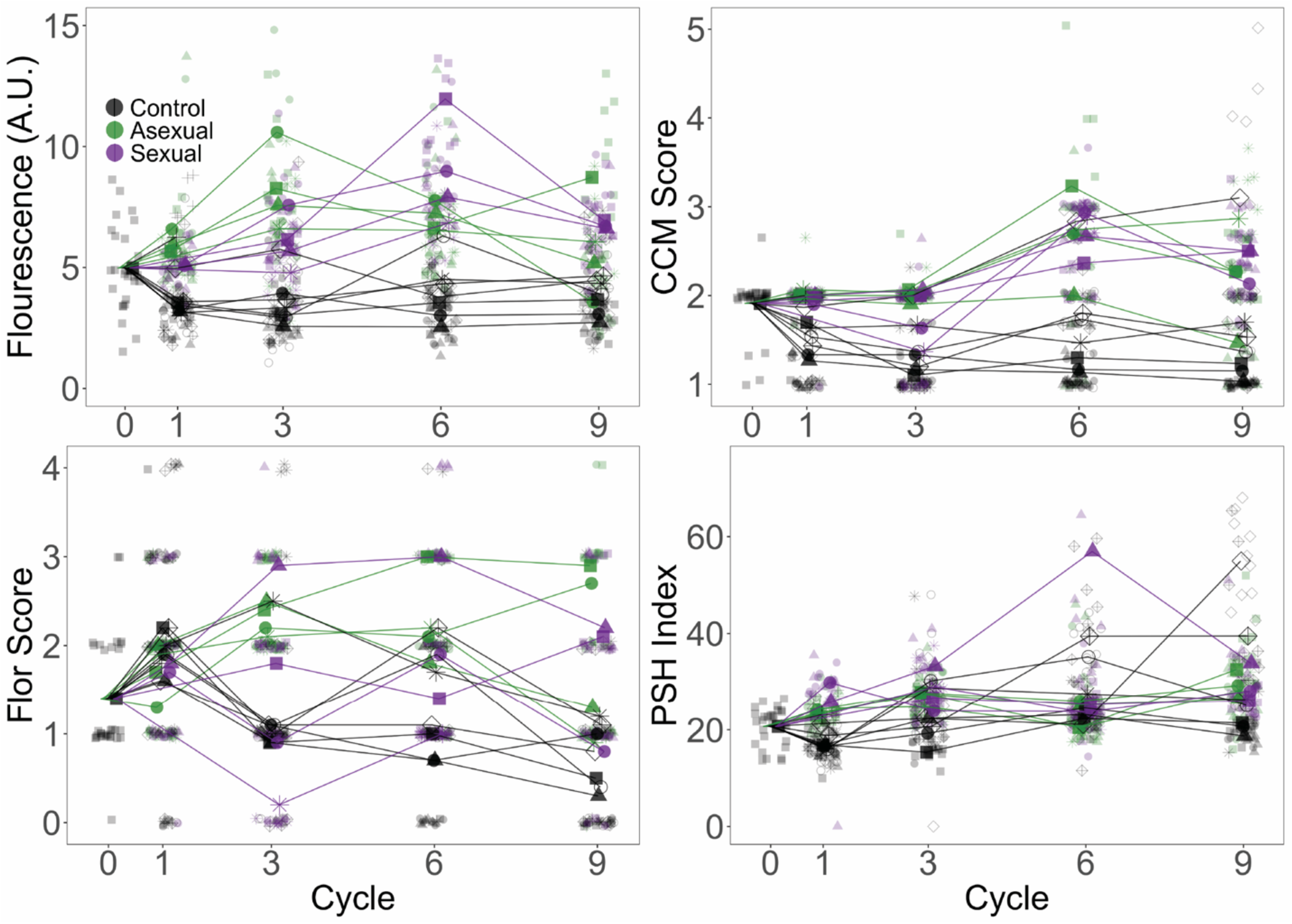
Evolved Phenotypes by Replicate Population for the YJM128 Background. Phenotypes were assessed as in Figure 2. Larger points represent replicate population averages, with each replicate represented by a different shape/color combination (as in Figure 4); smaller points represent measurements from individual clones. Fluorescence, CCM, and PSH are average values of three replicate measurements per clone; Flor is based on one replicate.

**Figure S4:**
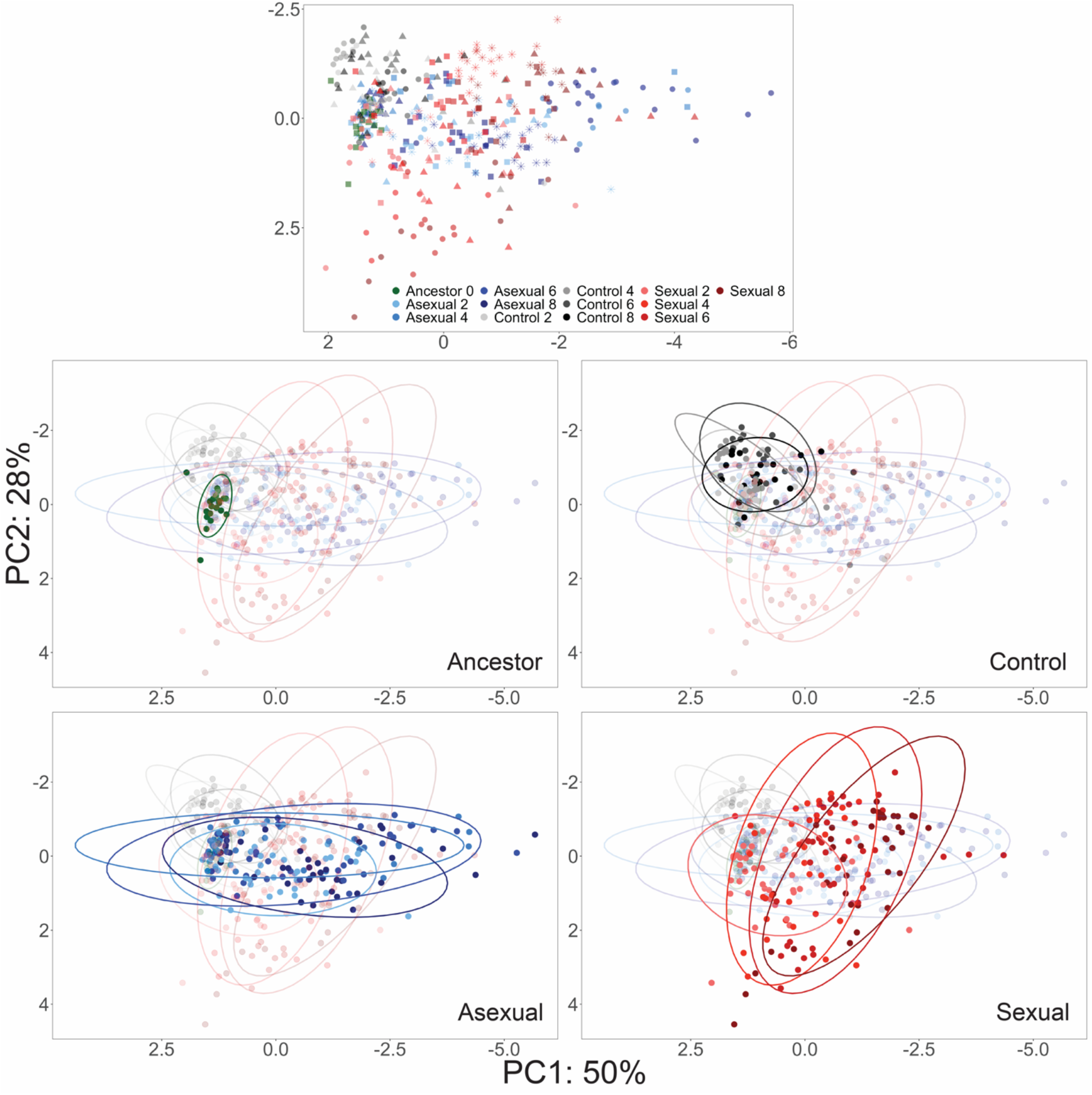
PCA of YJM311 Evolved Populations. Each point represents a single clone; ancestor-green, control-gray through black, asexual-light blue through dark blue, sexual-light red through dark red. Colors become darker as cycle number increases. In the first panel, shapes as in Figure 4 to highlight different replicate populations.

**Figure S5:**
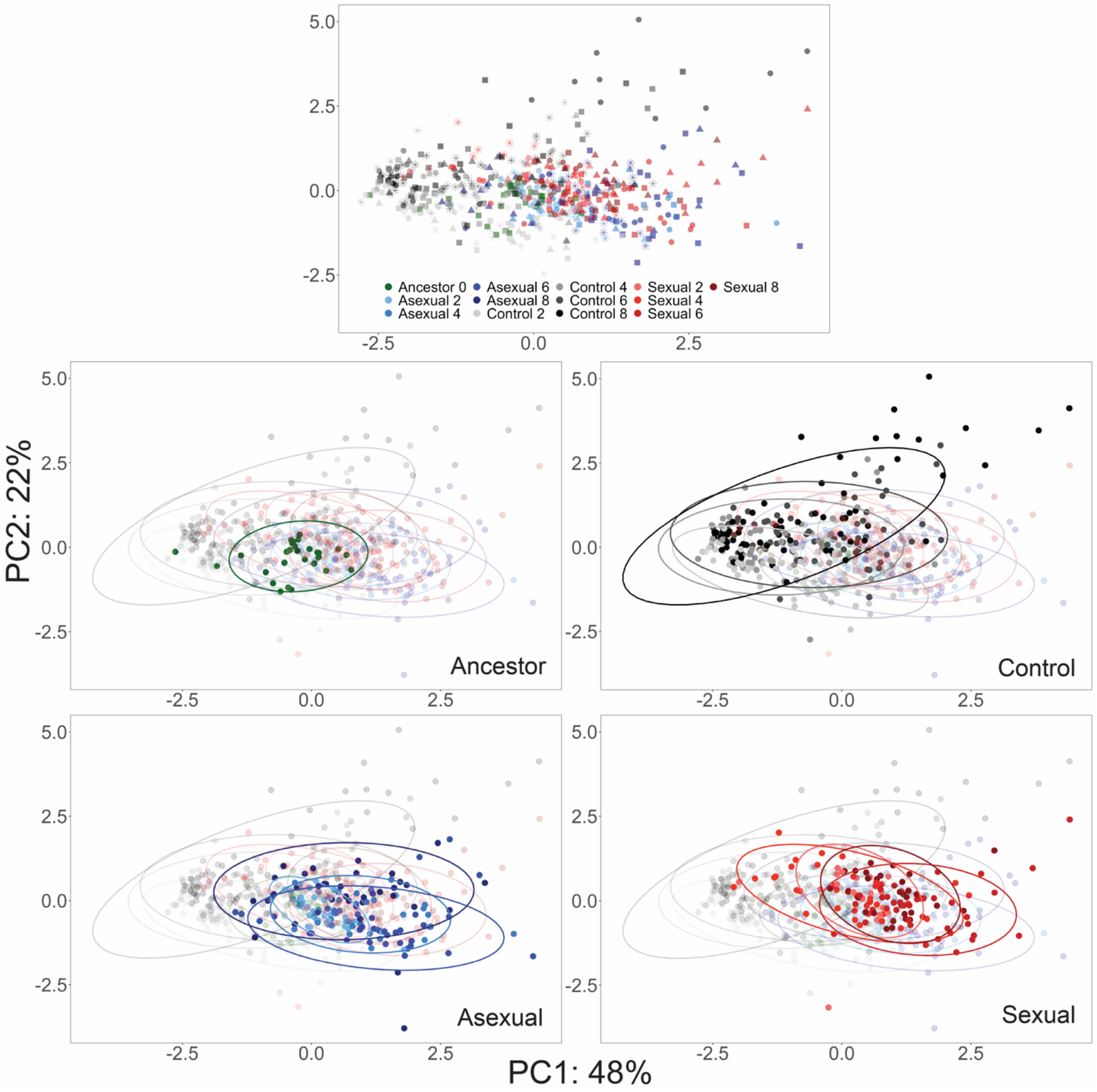
PCA of YJM128 Evolved Populations. Each point represents a single clone; ancestor-green, control-gray through black, asexual-light blue through dark blue, sexual-light red through dark red. Colors become darker as cycle number increases. In the first panel, shapes as in Figure 4 to highlight different replicate populations.

**Figure S6:**
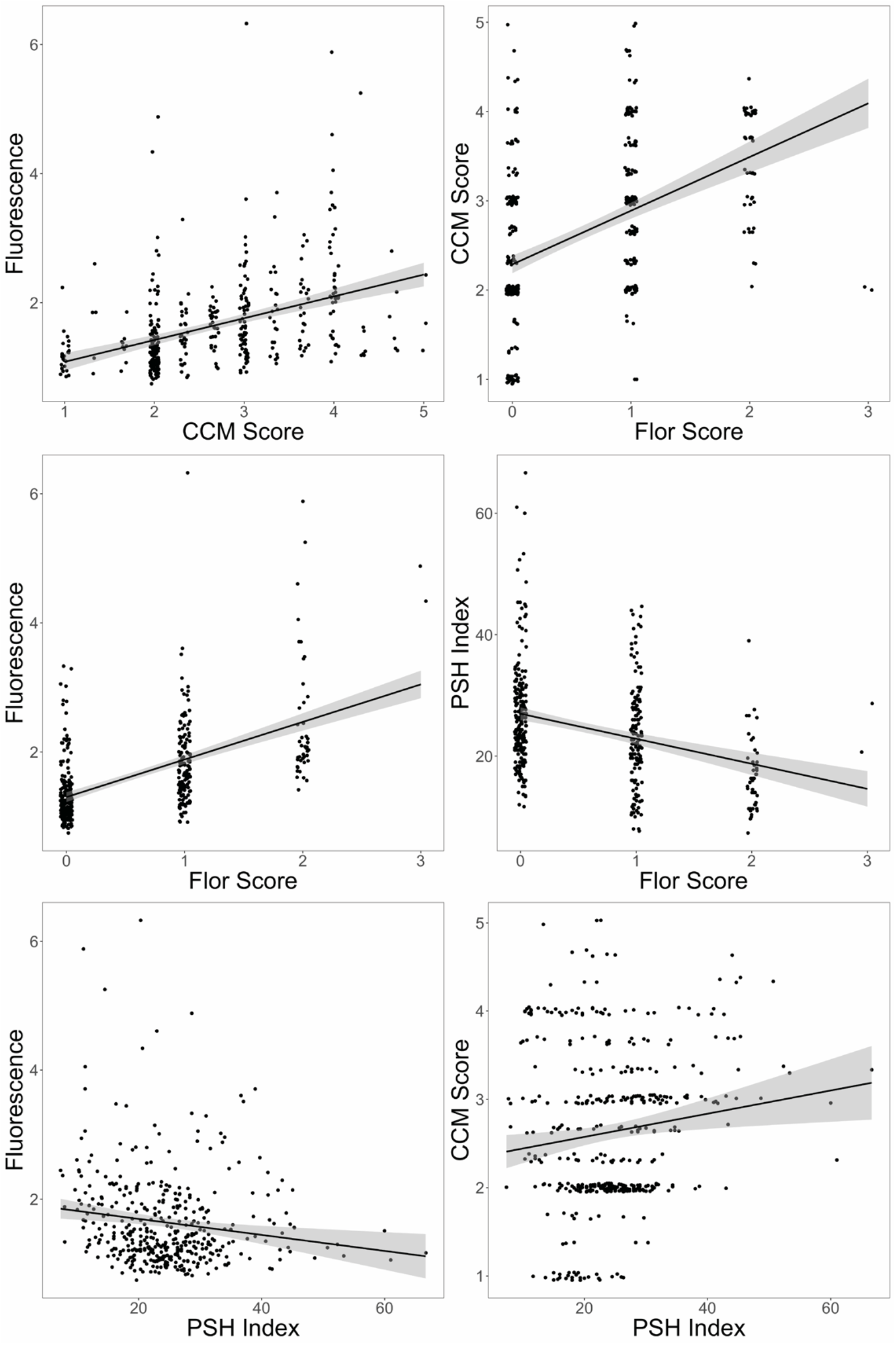
Correlations among phenotypes for individual clones from YJM311 populations. All correlations were significant with *p* < 0.001. The adjusted R^2^ for the relationships are as follows: PA vs. CCM: 0.167, PA vs. Flor: 0.306, PA vs. PSH: 0.020, CCM vs. Flor: 0.220, PSH vs Flor: 0.105, CCM vs. PSH: 0.015. Despite the statistical significance, most of these correlations explain little of the variance in the data; the presence of one multicellular phenotype does not necessarily have predict the presence of another.

**Figure S7:**
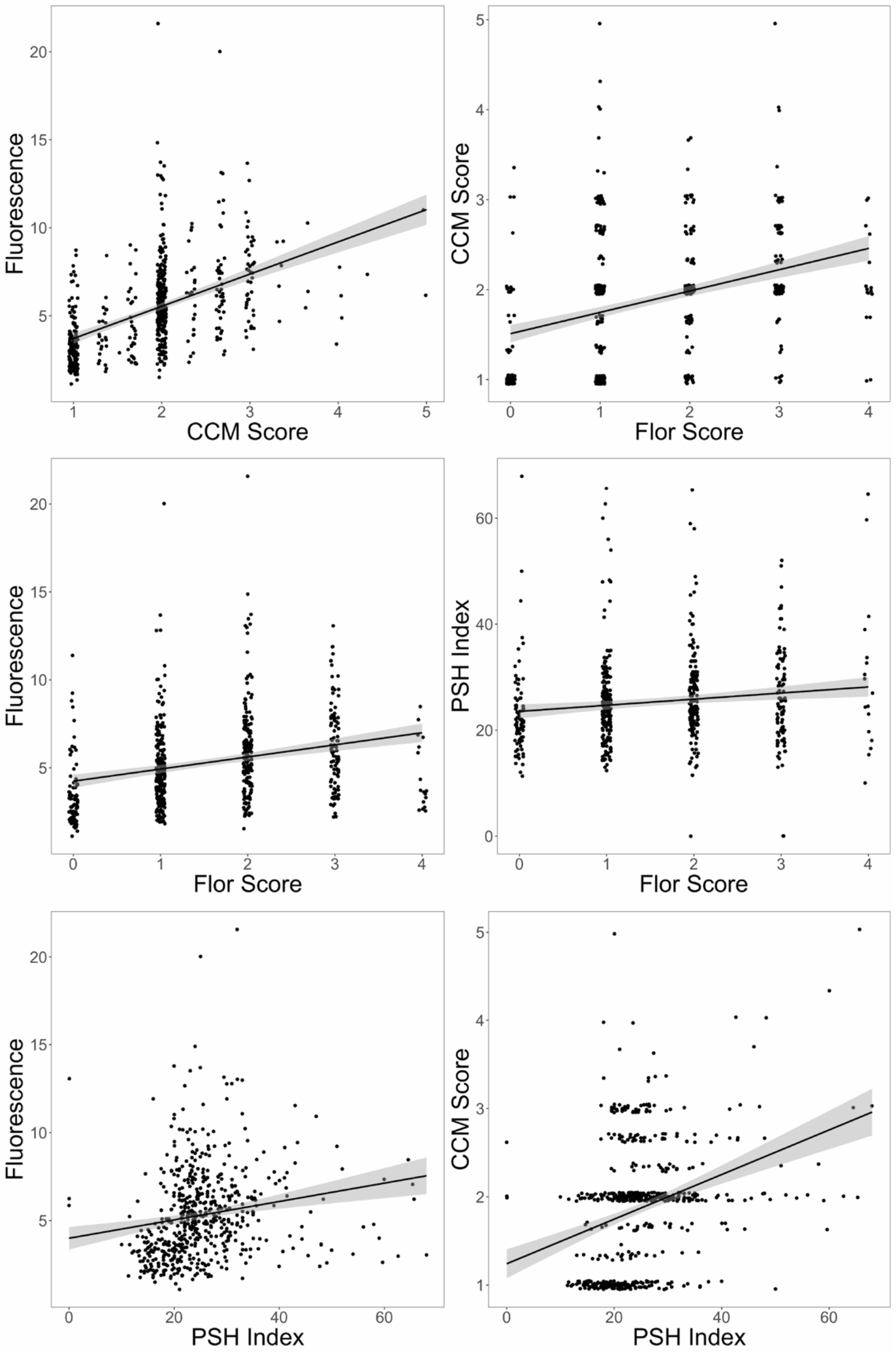
Correlations among phenotypes for individual clones from YJM128 populations. All correlations were significant with *p* < 0.001. The adjusted R^2^ for the relationships are as follows: PA vs. CCM: 0.235, PA vs. Flor: 0.071, PA vs. PSH: 0.0294, CCM vs. Flor: 0.121, PSH vs Flor: 0.0162, CCM vs. PSH: 0.102. Despite the statistical significance, most of these correlations explain little of the variance in the data; the presence of one multicellular phenotype does not necessarily predict the presence of another.

**Figure S8:**
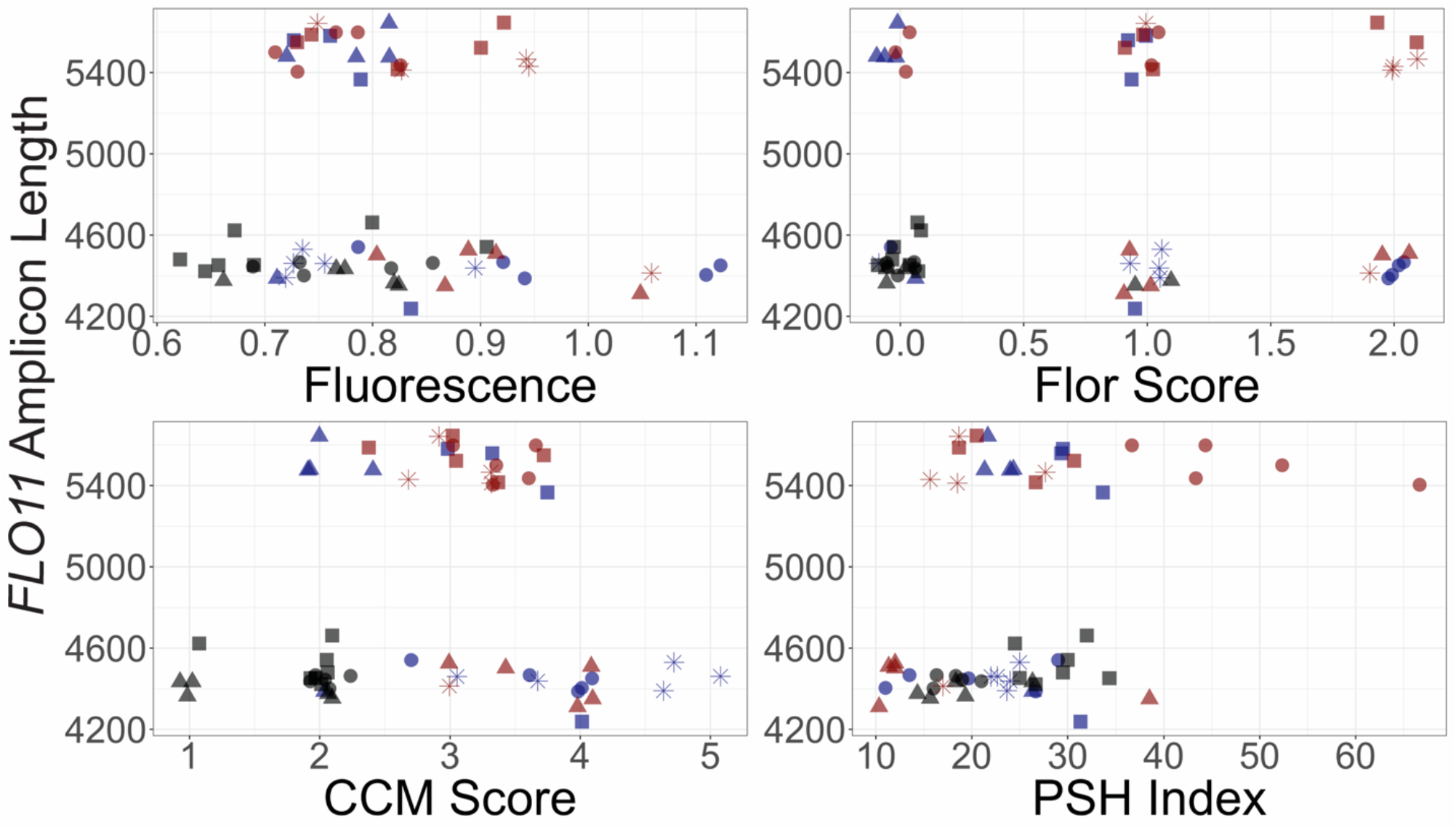
*FLO11* Length by Phenotype for the YJM311 Background. *FLO11* length measurements, point shapes and point colors as in Figure 3. Each panel plots the allele length for a clone with one of its phenotypic measurements.

**Figure S9:**
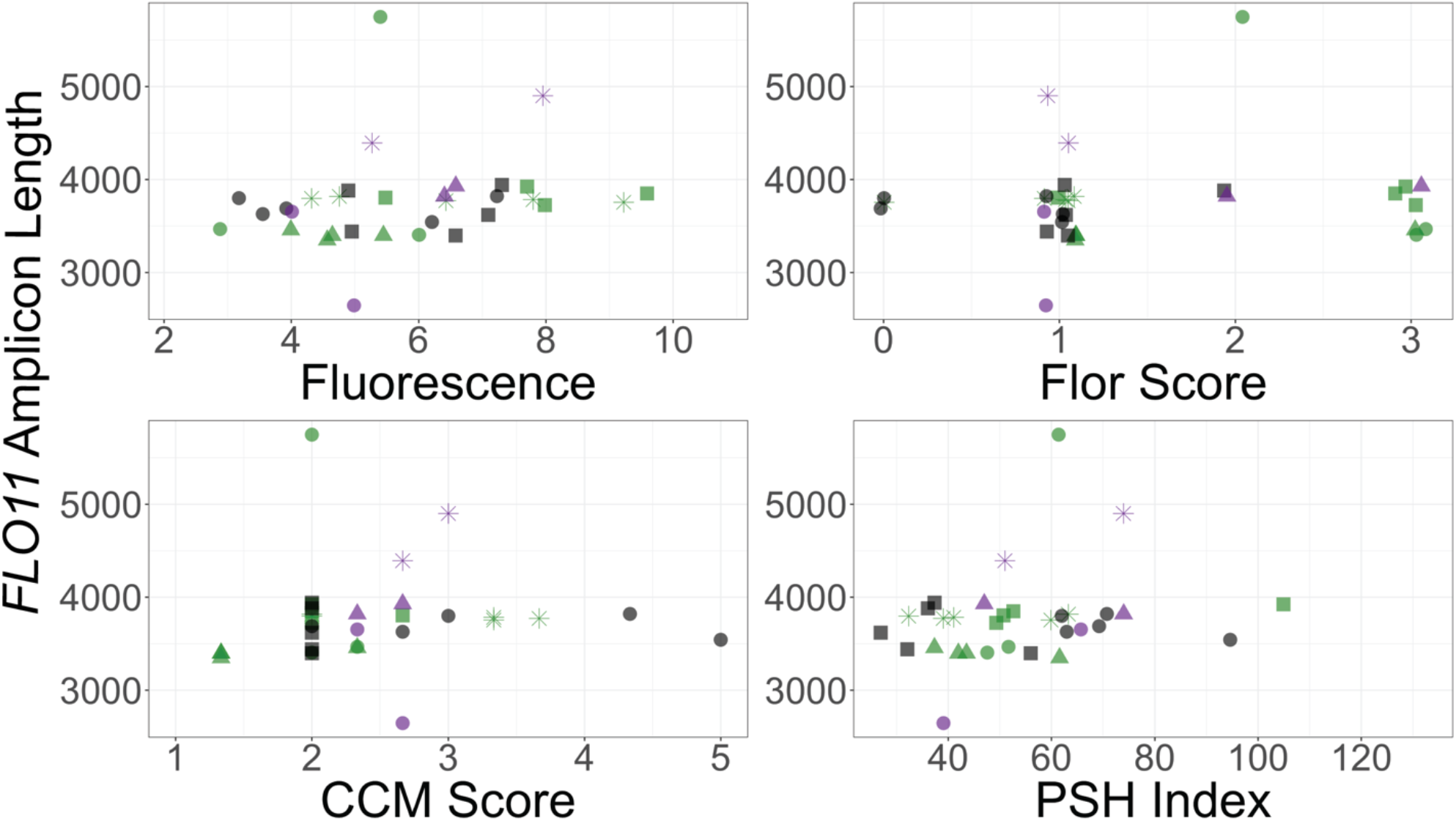
*FLO11* Length by Phenotype for the YJM128 Background. *FLO11* length measurements, point shapes and point colors as in Figure 3. Each panel plots the allele length for a clone with one of its phenotypic measurements.

**Figure S10:**
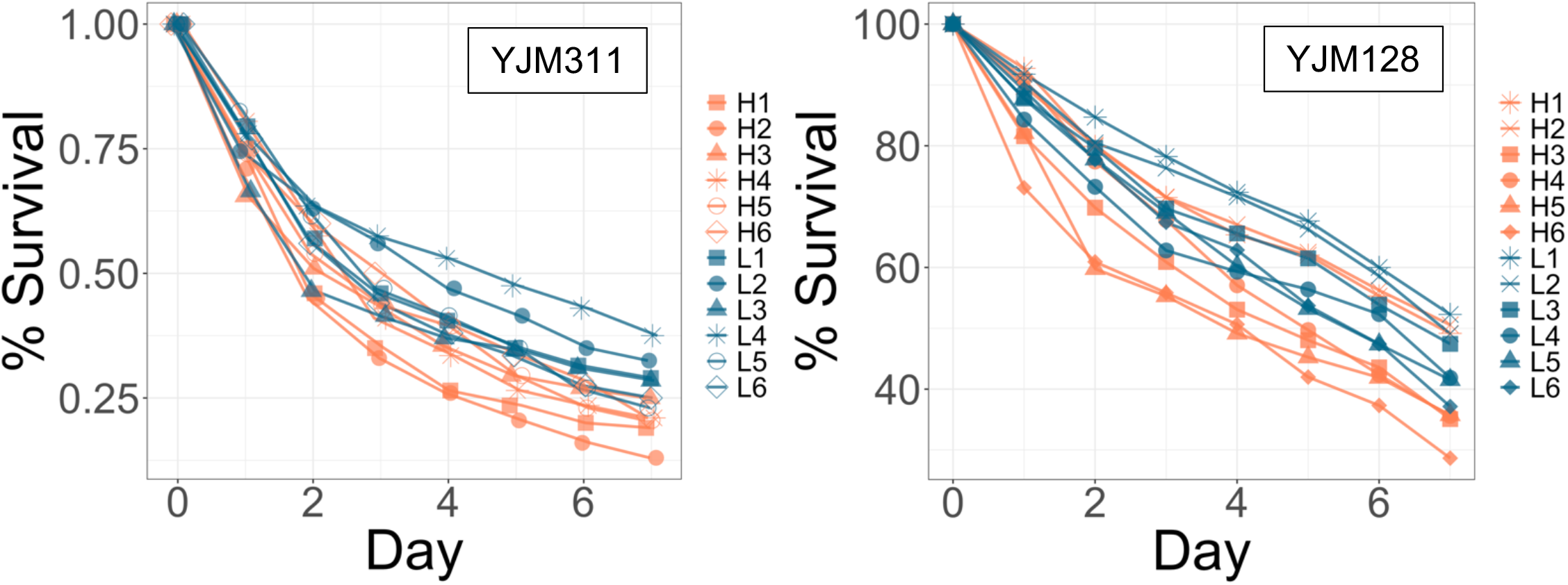
Survival Assay by Strain. Data as in Figure 4. Information on individual strains can be found in Tables S1 and S2.

